# The structural mechanism of MCIA complex assembly links mitochondrial redox pathways

**DOI:** 10.1101/2023.02.23.529646

**Authors:** Lindsay McGregor, Samira Acajjaoui, Ambroise Desfosses, Melissa Saïdi, Maria Bacia-Verloop, Jennifer J. Schwarz, Pauline Juyoux, Jill von Velsen, Matthew W. Bowler, Andrew McCarthy, Eaazhisai Kandiah, Gordon Leonard, Irina Gutsche, Montserrat Soler-Lopez

## Abstract

The mitochondrial Complex I assembly (MCIA) complex is an essential player in the biogenesis of respiratory Complex I (CI), the multiprotein complex responsible for the initiation of oxidative phosphorylation (OXPHOS). It is not well understood how MCIA facilitates the assembly of CI. Here we report the structural basis of the complex formation between the MCIA subunits ECSIT and ACAD9. ECSIT binding induces a major conformational change in the FAD-binding loop of ACAD9, resulting in efflux of the FAD cofactor and redeployment of ACAD9 from fatty acid β-oxidation (FAO) to CI assembly. We identify an adjacent α-helix as a key structural element that specifically enables the CI assembly functionality of ACAD9, distinguishing it from its closely related VLCAD counterpart. Furthermore, we show that ECSIT is phosphorylated *in vitro* and *ex cellulo* and provide evidence that phosphorylation downregulates its association with ACAD9. Interestingly, ECSIT has previously been linked to the pathogenesis of Alzheimer’s disease and here we show that ECSIT phosphorylation in neuronal cells is reduced upon exposure to amyloid-β (Aβ) oligomers.

These findings shed light on the assembly of the MCIA complex and implicate ECSIT as a potential reprogrammer of bioenergetic metabolic pathways that can be altered when mitochondria are affected by Aβ toxicity, a hallmark of Alzheimer’s disease.

## INTRODUCTION

A worldwide research effort is currently underway to identify the factors driving Alzheimer’s disease (AD) pathogenesis in order to find better ways to diagnose the disease, delay its onset and prevent progression. Due to their high energy demands and limited glycolysis rate, neurons depend on an efficient oxidative metabolism – *i.e*., oxidative phosphorylation (OXPHOS) and fatty acid beta-oxidation (FAO) – and are particularly vulnerable to mitochondrial dysfunctions. Because of this, age-related degradation of mitochondria is a prime suspect in AD pathophysiology^1^. Notably, in the brain of AD patients amyloid-β (Aβ) peptides progressively accumulate within mitochondria and perturb the mitochondrial respiratory Complex I (CI)^2^, a ∼1 MDa membrane protein complex composed of 44 different subunits (encoded in the mitochondrial and nuclear genome) that is essential for OXPHOS^3^. Despite its central importance, how CI is altered by Aβ and relates to neuronal integrity remains an open question.

While the molecular structure of the core CI subunits has been determined in atomic detail^4^, much less is known about the biogenesis of CI. This multistep process involves transiently associated assembly factors that integrate core and accessory/supernumerary subunits as well as cofactors into the final holoenzyme. A key player in CI biogenesis is the mitochondrial CI assembly (MCIA) complex^5^. This complex consists of three core proteins - NDUFAF1, ACAD9 and ECSIT - that appear to further associate with three peripheral membrane proteins^5^. The organisation of the MCIA complex and its role in CI assembly are unclear, partly because the individual MCIA components also mediate other cell functions. In particular, ACAD9 was annotated as an acyl-CoA dehydrogenase (ACAD) enzyme due to its sequence homology (47% amino acid identity and 65% amino acid similarity) with the very long chain acyl-CoA dehydrogenase VLCAD^6^, which both can initiate the FAO pathway with the concurrent reduction of their FAD cofactor^7^. In turn, ECSIT participates in cytoplasmic and nuclear signalling pathways, in which different ECSIT isoforms and post-translational modifications might contribute to its functional complexity^8^. Furthermore, ECSIT was identified as a molecular node interacting with Aβ-producing enzymes^9^, potentially implicating it in AD pathogenesis^10^.

We and others recently discovered that ECSIT binding triggers ACAD9 deflavination, switching it from an FAO enzyme to a CI assembly factor^11, 12^. These two mutually exclusive functions allow for the coordinated regulation between distinct multifunctional protein complexes to ensure efficient energy production. Here we provide high resolution structural insights into the MCIA subcomplex, ACAD9-ECSIT_CTER_, and identify the conformational changes that ACAD9 undergoes upon ECSIT binding. We provide experimental evidence of *in vitro* and *ex cellulo* ECSIT phosphorylation and its effect on the ACAD9-ECSIT interaction and reveal a decrease in ECSIT phosphorylation levels upon exposure to Aβ oligomer toxicity. These findings will pave the way for evaluating MCIA proteins as potential diagnostic biomarkers for detecting early AD pre-symptomatic stages when mitochondria are primarily affected by Aβ toxicity^13^.

## RESULTS

### Cryo-EM structure of the ACAD9-ECSIT_CTER_ complex

We previously reported a cryo-electron microscopy (cryo-EM) structure of ACAD9 in complex with a C-terminal fragment of ECSIT (ECSIT_CTD_, residues 247-431) at low (∼15 Å) resolution^11^. The inclusion of an additional N-terminal 26 residues in the ECSIT construct (hereafter named ECSIT_CTER_, residues 221-431), combined with optimisation of the purification and subcomplex reconstitution protocols (**Fig. S1A-D**) led to a 3.0 Å resolution cryo-EM structure of the ACAD9-ECSIT_CTER_ complex (**Fig. 1, Figs. S2, Table S1**), enabling us to build and refine an atomic model that includes nearly all ACAD9 residues as well as a 15-residue peptide spanning ECSIT residues 320-334 (**Fig. 1A-B, Fig. S3A**). As previously described^14^, ACAD9 is present as a dimer, where the C-terminal regions (residues 600-621) form helix loops that are positioned around the neighbouring protomer thereby stabilising the active dimer (**Fig 1B**). Such helix swapping has been suggested to promote the stability of the homodimer in structural homologues adopting the same C2-symmetry^15^. The ACAD9 monomer consists of an N-terminal dehydrogenase domain (res. 38-453 comprising an α-helical, a β-sheet and a second α-helical subdomains) and a C-terminal α-helical bundle vestigial domain (res. 487-587) connected by a poorly ordered ∼35-residue linker (**Fig. 1C**). As expected, no significant density was observed for the flavin adenine dinucleotide (FAD) in the dehydrogenase domain, confirming our previous observation that the binding of ECSIT triggers ejection of the cofactor from ACAD9^11^.

**Figure 1:**
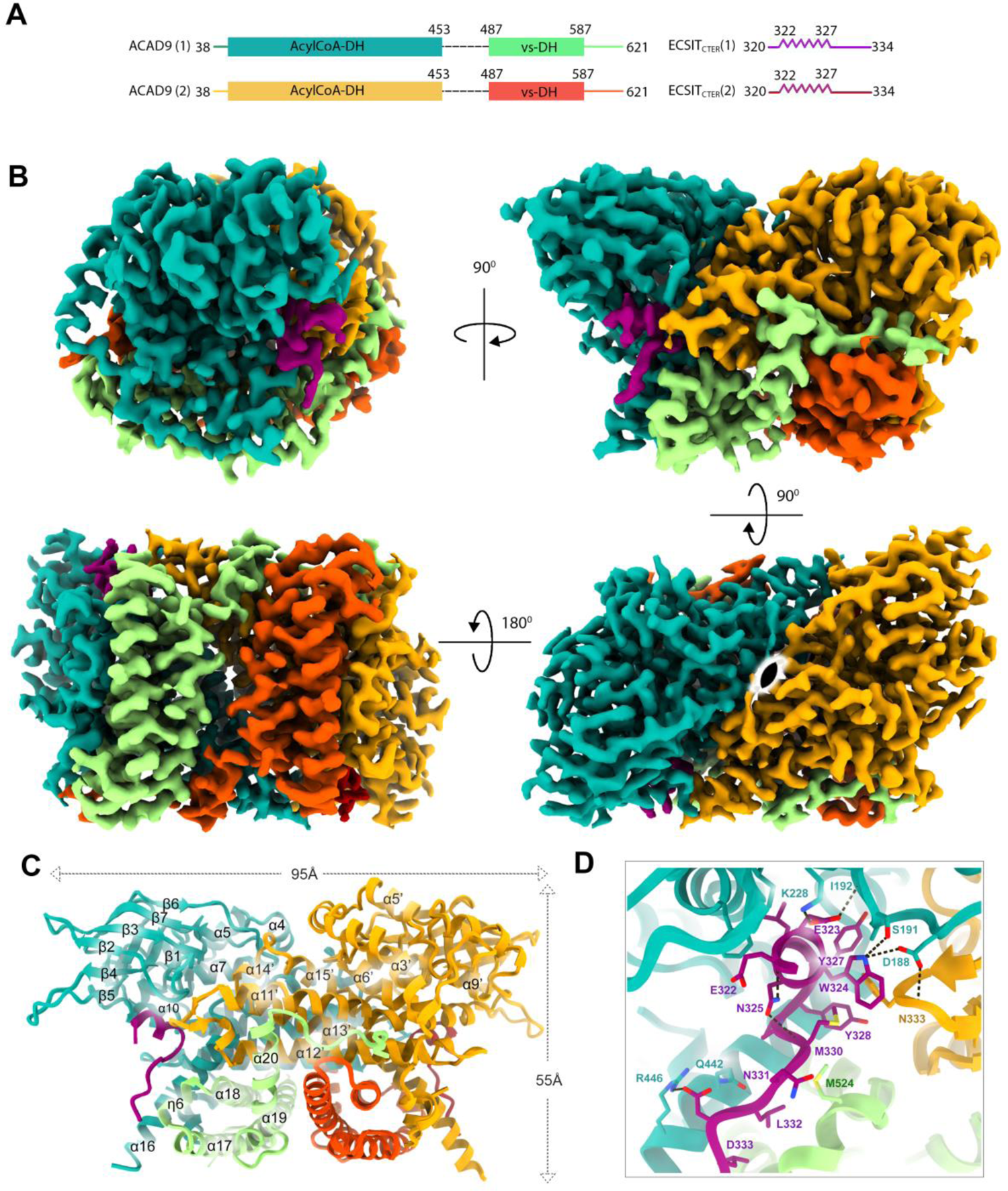
Cryo-EM structure of the ACAD9-ECSIT_CTER_ complex. **(A)** Schematic of the sequences of ACAD9 and ECSIT_CTER_ modelled in the cryo-EM map. The two ACAD9 monomers are labelled (1) and (2), with their dehydrogenase domains shown in teal and light orange, and their vestigial domains in light green and dark orange, respectively. ECSIT_CTER_ monomers are in purple and red. The dashed lines indicate ACAD9 residues 454-486 absent from the final model. **(B)** Cryo-EM map of ACAD9-ECSIT_CTER_ showing side, front, top and bottom views (clockwise from top left) coloured domain-wise as in (A), with C2 symmetry represented (bottom right). Dimensions in width and height are indicated in Å**. (C)** Ribbon diagram of the refined structure of the ACAD9-ECSIT_CTER_ complex coloured as in A. **(D)** Key interactions involved in ECSIT_CTER_ recognition by the ACAD9 dehydrogenase/vestigial interface. Specifically, ECSIT Glu323 makes a salt bridge with ACAD9 Lys228, stabilising the ECSIT binding on the N-terminal segment with the ACAD9 β4 strand, whereas a second salt bridge between ECSIT Asp333 and ACAD9 Arg446 stabilises the ECSIT binding on the C-terminal end with the ACAD9 α16 helix. Furthermore, ECSIT directly interacts with the ACAD9 β1-β2 loop through three hydrogen bonds between ECSIT Trp324 and ACAD9 Ser191 and Asp188, respectively, and ECSIT Glu323 with the backbone amide of Ile192. Notably, while there is no evidence of direct ECSIT interactions with the second ACAD9 protomer, Asp188 shows an additional hydrogen bond with Asn333 on α11’-α12’ loop, bridging ECSIT binding with the ACAD9 dimer. Numerous intramolecular hydrogen bonds within ECSIT contribute to the stability of the conformation. In particular, Thr320 forms a hydrogen bond with Gln331 while Asn325 interacts with the backbone amide of Glu322 and Met330, respectively.

The ECSIT binding site is located at the junction of the ACAD9 dehydrogenase and vestigial domains (**Fig. 1A-D**), with one ECSIT_CTER_ per ACAD9 protomer. ECSIT residues 322-327 form a 3_10_-helix that interacts with the ACAD9 β1-β2 loop, while ECSIT residues 328-334 adopt an extended loop conformation lying in anti-parallel orientation between the ACAD9 α16 and η6 helices (**Fig. 1C-D**). Examination of the electrostatics properties of the binding interface shows that the surface of the ACAD9 is mainly positively charged whereas the modelled ECSIT sequence is mostly negative (**Fig. S3B**). Details of ECSIT_CTER_ recognition by the ACAD9 dehydrogenase/vestigial interface, showing key binding interactions, are summarized in **Fig. 1D**.

The relevance of the hydrogen bond between ECSIT Trp324 and ACAD9 Ser191 (**Fig. 1D**) was confirmed by a reduction in complex formation of ACAD9 with an ECSIT-W324A mutant observed by mass photometry and DLS (**Fig. 2B, G**). Conversely, an ACAD9-S191A mutant showed higher stability than ACAD9_WT_, with less dissociation into monomers (**Fig. S1B, Fig. 2A**), and retained the ability to bind ECSIT_CTER_ (**Fig. 2A, G**) even if showing a lower affinity by ITC (**Fig. S1D, Fig. 2H**). Interestingly, despite their positioning in the ACAD9 binding pocket (**Fig. 1D-E**), ECSIT residues Tyr327 and Tyr328 do not appear to strongly interact with any ACAD9 residues (**Fig. 1D**). Replacement of these Tyrosine residues by either Phenylalanines or Alanines shows that the loss of the hydrophobic nature/π-bonding and hydrogen bonding capabilities of Tyr327 affects complex formation more than removal of the hydrogen bonding partner only, whereas removal of both the hydrophobic/π-bonding and hydrogen bonding properties of Tyr328 permitted complex formation (**Fig. 2C-G**).

**Figure 2:**
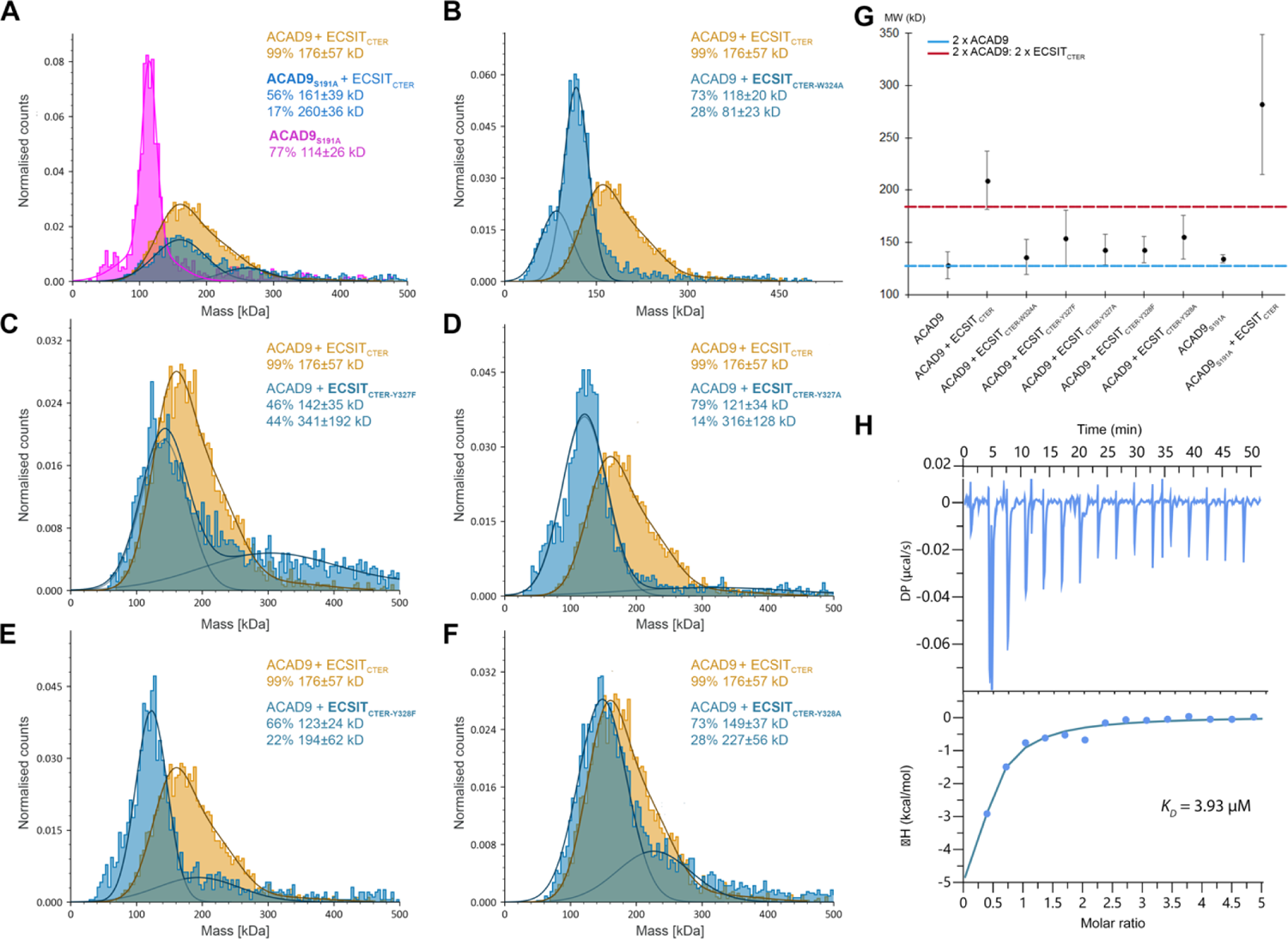
Analysis of interface derived mutants on ACAD9-ECSIT complex formation. A range of mutants were designed to probe the complex interface and identify the role of the residues in complex formation. The mutant species were reconstituted with the corresponding WT protein and compared with ACAD9_WT_-ECSIT_CTER_. **(A)** ACAD9_S191A_ appears a more stable dimer than ACAD9_WT_ (Fig. S1B) and forms an ACAD9_S191A_-ECSIT_CTER_ complex. **(B)** ECSIT_CTER-W324A_ seems to negatively impact complex formation, with the MW of the main species similar to ACAD9_WT_. **(C)** ECSIT_CTER-Y327F_ complexes with ACAD9_WT_ but also forms higher order species. **(D)** ECSIT_CTER-Y327A_ affects complex formation, with the MW of the main species similar to ACAD9_WT_. **(E)** ECSIT_CTER-Y328F_ has a detrimental effect on complex formation, whereby the MW of the main species is similar to ACAD9_WT_ whereas **(F)** ECSIT_CTER-Y328A_forms the complex, however, with additional higher MW species. **(G)** DLS was used to evaluate the MW of the mutants in complex with the corresponding WT partner. All ECSIT mutants show a reduced average MW (130 – 150 kDa) in comparison to the ACAD9-ECSIT_CTER_ complex, possibly indicating partial complex formation but a reduction in stability. ACAD9_S191A_ has a MW very similar to ACAD9_WT_, however, in complex with ECSIT_CTER_, there is the formation of higher MW species. **(H)** ITC binding assay for the binding affinity between ACAD9_S191A_ and ECSIT_CTER_. The equilibrium dissociation constant (K_D_) of the ACAD9_S191A_-ECSIT_CTER_ complex is 3.93 μM, ∼ 3-fold lower than that of ACAD9_WT_-ECSIT_CTER_ (Fig. S1D).

For additional insights, we used AlphaFold2 (AF2)^16^ to predict the structure of the ACAD9-ECSIT complex using sequences matching our constructs. The AF2 model predicted the binding site to contain ECSIT residues 316-338 and closely resembled the experimental structure (RMSD of 1.635 Å over 552 Cα atoms; **Fig. S4A-B, D**). No additional ECSIT residues were predicted to contact ACAD9, suggesting that most, if not all, of the specific interactions between these proteins were captured in the cryo-EM structure. Indeed, we found that a synthetic peptide spanning ECSIT residues 318-336 was sufficient to eject FAD from ACAD9 (**Fig. S1E**), confirming that these residues are crucial for complex formation. Notably, the full length AF2 model predicts that ECSIT comprises two flexibly linked globular domains, an N-terminal domain (res. 70-208) with a pentatricopeptide repeat fold and a C-terminal domain (res. 221-392) with an RNAseH-like fold comprising a six-stranded mixed β-sheet and four short helices^17^. The ACAD9-binding residues of ECSIT reside within the C-terminal domain and localise to the tip of a long loop (res. 306-357) flexibly connected to strands β3 and β4 of the RNAseH-like domain. This turns to be very close to a density extension observed in our earlier low-resolution map of ACAD9-ECSIT_CTD_ ^11^ (**Fig. S4C**), allowing this domain to be positioned within the extra density with only a minor adjustment of the long predicted β4-β5 loop. The long linkers (residues 306-320 and 335-356) (**Fig. S4A-B**) connecting the ACAD9 binding region of ECSIT to the RNAseH-like C-terminal domain likely contribute to the high flexibility observed in biophysical analyses^11^ (**Fig. 1A, Fig. S4A**).

### Cryo-EM structure of ACAD9 reveals the opening of a gatekeeper loop upon ECSIT binding

In order to identify structural changes that occur in ACAD9 upon ECSIT binding, we pursued a cryo-EM structural study of ACAD9_WT_ in the unbound state. Given a strongly preferred orientation (**Fig. S5A-D**) resulting in an anisotropic 3D reconstruction and limiting the resolution of the final map to ∼5.0 – 5.5 Å in the core and ∼ 6.5 – 8.0 Å in the periphery (**Fig. S5D**), the analysis was restricted to a rigid body fit of the ACAD9 AF2 model (residues 38-621). Considering the observed stability, compactness and ECSIT-binding ability of the ACAD9_S191A_ mutant (**Fig. 2A, G**), we chose it as a suitable candidate for further improving the ACAD9 structure. To increase the orientational diversity, we collected a 35° tilted cryo-EM data set (**Fig. S5E-H**) and solved the structure of ACAD9_S191A_ at 3.6 Å resolution (**Fig. S6A-C**). Consistent with FAO activity, the structure features one molecule of FAD in the cofactor binding site of each ACAD9 protomer (**Fig. S6C, D**). As expected, no significant density occupies the ECSIT binding site identified in the ACAD9_WT_-ECSIT_CTER_ structure.

Strikingly, comparing the structures of ACAD9 alone and in complex with ECSIT_CTER_ reveals that, in addition to causing FAD ejection, ECSIT binding induces a large conformational change in the β1-β2 loop adjacent to the FAD binding site (**Fig. 3**). In particular, the structure of ACAD9_S191A_ shows that this loop adopts a closed, downward facing position, thereby acting as a barrier between the adenine nucleotide of the FAD molecule and the internal core of ACAD9 (**Fig. 3A, C**). In contrast, the loop adopts a very different conformation in the ACAD9_WT_-ECSIT_CTER_ model (**Fig. 3B, D, E**), where the tip of the loop (residue Gly186) moves by ∼10 Å upwards to allow the ECSIT 3_10_-helix to insert, suggesting that this mobile ACAD9 β1-β2 loop plays the role of a gatekeeper (**Fig. 3F**). Although it does not directly interact with the FAD molecule, we suggest that removal of this barrier destabilises the cofactor environment, leading to its ejection and the reassignment of ACAD9 from an FAO enzyme to a CI assembly factor. The downward facing loop position in ACAD9_S191A_ clashes with the superposed ECSIT 3_10_-helix indicating that the flipping mechanism is essential for ECSIT binding and MCIA formation.

**Figure 3:**
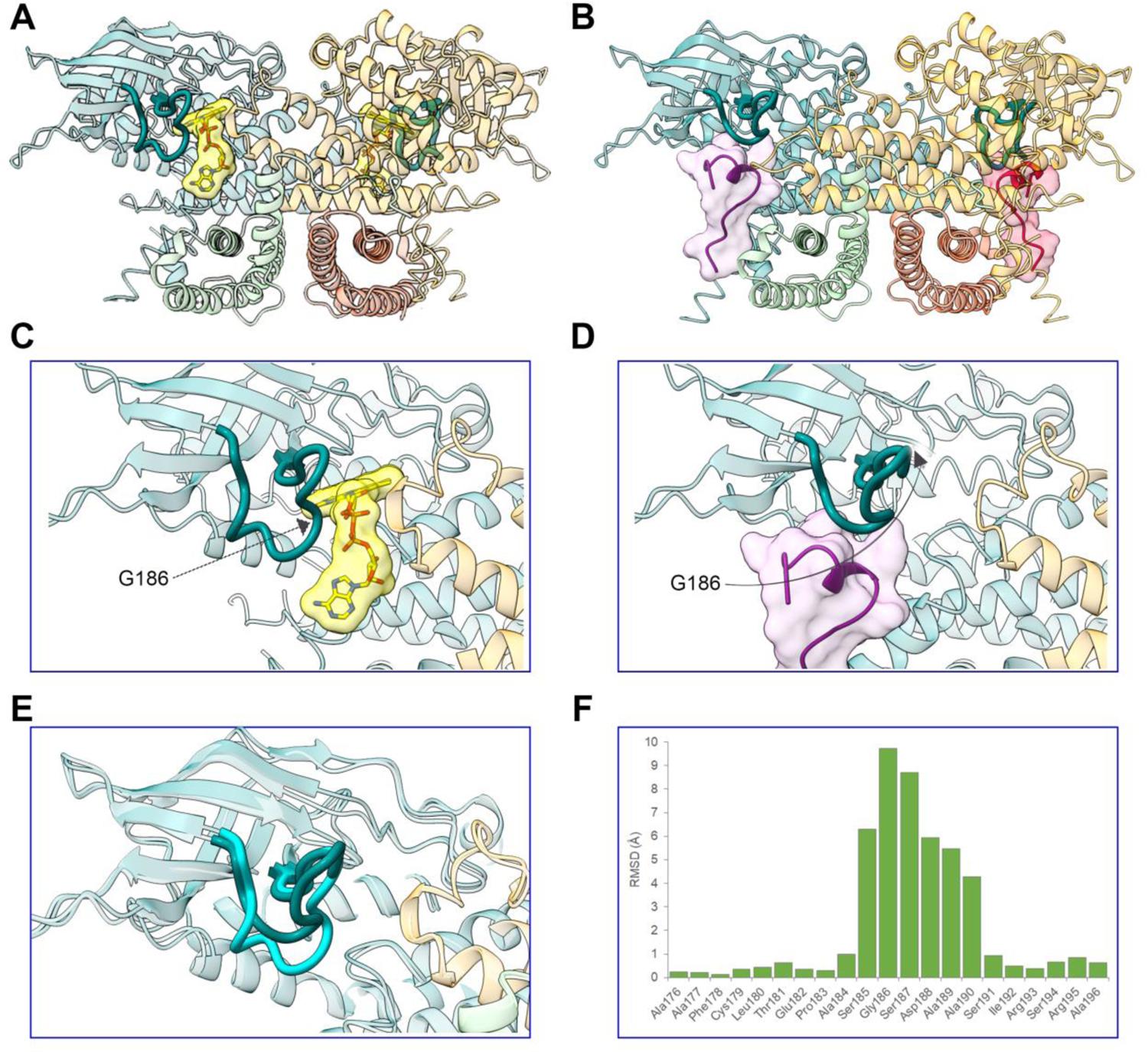
A gatekeeper loop movement of 10 Å is induced upon ECSIT_CTER_ binding to ACAD9. ECSIT binding to ACAD9 induces the ejection of the FAD molecule and the displacement of a loop bridging the ECSIT:FAD binding sites. **(A)** ACAD9_S191A_ in dehydrogenase form, with the bound FAD highlighted in yellow. The β1-β2 loop consisting of ACAD9 residues 178-195 is depicted in dark teal. **(B)** ACAD9-ECSIT_CTER_ complex showing the location of the two ECSIT binding sites adjacent to ACAD9 residues F178-R195 in bold. **(C)** Close-up of (A), showing the loop adopting a downfacing position, in a ‘closed conformation’, acting as a barrier to the internal core of ACAD9 and the FAD pocket. **(D)** Close-up of (B), after ECSIT binds to ACAD9. Residue Gly186 is on the tip of the β1-β2 loop and is displaced by ∼10 Å, flipping upwards into an ‘open conformation’. **(E)** Superposition of the β1-β2 loop in ACAD9 unbound (cyan) and in complex with ECSIT_CTER_ (dark teal). **(F)** RMSD of Cα positions of the β1-β2 loop residues in ACAD9 unbound vs. in complex with ECSIT_CTER_.

Similarly to the ACAD9_WT_-ECSIT_CTER_ dataset, residues 455-485 could not be built into the ACAD9_S191A_ map. However, increasing the threshold of the unsharpened ACAD9_WT_ map reveals some density at the base of ACAD9_S191A_, hinting at their possible position. Thus, we fitted the missing residues into this signal, using the coordinates of the ACAD9 AF2 model where residues 455-464 and 475-492 form flexible loops, sandwiching a short α-helix (res. 465-474). The lack of definition in the density points to a mobility of this region (**Fig. S6E**) and may be explained by the lack of contacts between the α-helix and the ACAD9 core. Interestingly, we do not see signal for these residues in the ACAD9_S191A_ dataset, probably due to a larger number of particles being included in the higher resolution reconstruction, leading to the signal being averaged out due to high mobility.

### Cryo-EM structure of ACAD9 reveals unique features in comparison to other ACADs

The comparison of our ACAD9_S191A_ structure with both the VLCAD-based ACAD9 homology model^6^ and a very recently published VLCAD crystal structure (PDB:7S7G ^18^) revealed a high structural similarity, with RMSD values of 1.16 Å and 1.11 Å, respectively (**Fig. S7A**). A visualisation of the evolutionary conserved residues also reveals that the dehydrogenase domains are quite conserved within the ACAD family^19^ (**Fig. S7B**). Intriguingly, and in agreement with previous studies^6,11^, ACAD9 is the only ACAD family member capable of binding ECSIT and assisting CI assembly (**Fig. S1D**), whereas VLCAD does not fulfil these functions^11^. Thus, given that the ECSIT binding induces the flipping of the gatekeeper loop, we generated ACAD9 point mutants intended to mimic the VLCAD sequence on this loop and analysed them by DLS and mass photometry. In regards to their ECSIT-binding properties, these mutants behave essentially like ACAD9_WT_ (**Figs. S7B-G**), suggesting that, although the conformational flexibility of the ACAD9 gatekeeper loop is important for the dimer stability and the enzymatic activity, it is nevertheless not decisive to confer its ability to bind ECSIT.

Therefore, in our quest to identify the distinct ECSIT binding features of ACAD9, we compared the FAD cofactor sites between our ACAD9 structure and the reported VLCAD structure (**Fig. S8A-D**)^15^. In ACAD9, the FAD molecule is situated on top of the core strands and adopts an elongated conformation for maximal interaction with the FAD-binding domain (**Fig. S6D, Fig. S8D**), very similar to the FAD positioning in VLCAD. The overall architecture of the pocket is analogous in both proteins. However, in ACAD9 the volume of the FAD binding pocket is larger and bonding interactions between the ACAD9 protein and the FAD molecule are longer and weaker, suggesting that the fold of the FAD-binding domain and the cofactor conformation are not independent, whereby the arrangement of the domain determines the position and shape of the pocket and thus the cofactor conformation.

Interestingly, in VLCAD, two Methionine residues located on a 3_10_-helix (res. 437-443) from the second protomer make sulphur-π interactions with a Tryptophan residue (W209) and the FAD isoalloxazine moiety of the first protomer, while in ACAD9 the corresponding Leucine and Threonine residues are unable to mediate similar interactions with the equivalent Tryptophan (W213) (**Fig. S8C-D**). We thus generated an ACAD9 mutant, ACAD9_VLCAD_, to mimic VLCAD by replacing residues ^401^LGGLGYT^407^ by the corresponding VLCAD residues ^437^MGGMGFM^4^^43^ (**Fig. S8F-G**). Although ACAD9_VLCAD_ forms a stable dimer, exhibiting a profile similar to VLCAD by mass photometry (**Fig. S8F**), it does not appear to form a complex with ECSIT_CTER_. Furthermore, the presence of ECSIT_CTER_ seems to induce dissociation of ACAD9_VLCAD_ into monomers (**Fig. S8F-G**). On the other hand, although the counterpart VLCAD_ACAD9_ mutant (**Fig. S8E, G**) appears to form a less stable dimer than the VLCAD_WT_, it forms a complex with ECSIT_CTER_, even if larger species also appear (**Fig. S8E, G**). Collectively, these findings suggest that in VLCAD, the dyad of Methionine residues located on a helix in the N-terminal α-dom2 of the neighbouring protomer (as described in ^15^) are responsible for stabilising the FAD in the binding pocket (**Fig. S8C-D**). This ability to retain the FAD cofactor is directly correlated with the stability of the homodimer but inversely correlated with the ability to bind ECSIT.

Lastly, another potential site that could account for the differences between ACAD9 and VLCAD in CI assembly lies on the 35-residue linker (res. 450-485 in ACAD9 and 486-521 in VLCAD), which shows the highest sequence divergence between the two proteins^6^. The recently reported VLCAD crystal structure models most of those residues (res. 486-499) facing away from the protein and projecting outward ^18^. This extended loop mediates *trans* interactions with the equivalent loop of another symmetry related VLCAD molecule (**Fig. S9A**), stabilising the quaternary conformation of a dimer of homodimers ^18^. However, superposition with the ACAD9_WT_ structure reveals that in ACAD9 this loop tucks into the core of the same homodimer instead (**Fig. S9B**). The higher conformational flexibility in VLCAD could be accounted by Gly488 at the end of α16 helix, which turns to be a hydrophobic residue (Ile452) in ACAD9 (**Fig. S9B**). Excitingly, alignment with the ACAD9-ECSIT_CTER_ complex structure shows that residues of the VLCAD flexible loop are in fact positioned near the ECSIT binding site in the ACAD9-ECSIT_CTER_ complex (**Fig. 1D, Fig. S9C**). Therefore, it appears that the equivalent ‘ECSIT binding’ site on VLCAD can be obstructed by the loop from another monomer. In fact, a structural alignment on this region between VLCAD residues 486-498 and ECSIT residues 320-334 (reversed to match VLCAD sequence positioning, 334-320) shows some overlap in hydrophobic character (**Fig. S9D**). In sum, these observations suggest that even if VLCAD contains an equivalent ‘ECSIT binding site’, it is unable to specifically bind ECSIT as it can be blocked by VLCAD itself. This auto-inhibitory feature would not be the case for ACAD9, as the equivalent sequence of residues are dissimilar (**Fig. S9D**), indicating another possible reason for the specificity of ACAD9 for ECSIT in comparison to VLCAD.

### ECSIT_CTER_ is phosphorylated and has an impact on MCIA complex stability

In addition to its pivotal role in the MCIA complex, ECSIT was also described as providing a bridge between Toll-like receptor (TLR) signal transduction and downstream signalling kinases^8^. Previous studies have shown evidence of ECSIT ubiquitination by the E3 ubiquitin ligase TRAF6 in the cytoplasm, which leads to ECSIT enrichment at the mitochondrial periphery to generate mROS^20^ and its translocation into the nucleus to regulate NF-κB activity^21^. Furthermore, another study reported a non-identified post-translationally modified form of ECSIT dependent on a MAPK kinase activity and apparently required for its role in the TLR pathway^8^. Therefore, to gain an insight into ECSIT regulatory mechanisms and given the frequent crosstalk between ubiquitination and phosphorylation, we investigated whether ECSIT undergoes phosphorylation in the presence of MAPK kinases and whether this has an impact on ACAD9-ECSIT subcomplex formation.

We first examined the potential phosphorylation of ECSIT *in vitro* by incubating ECSIT_CTER_ with a well-characterised MAPK kinase, p38α^22^. Denaturing gel electrophoresis showed that ECSIT_CTER_ migrates with higher mobility upon p38α kinase treatment (**Fig. 4A**), consistent with the gain of one or more negatively charged phosphate groups. Subsequent radiolabelling analysis confirmed the presence of strong ^32^P-labeled bands in ECSIT_CTER_ treated with activated p38α, indicating the direct phosphorylation of ECSIT by p38 MAPK kinase *in vitro* (**Fig. 4B**). Analysis of the faster migrating band by proteolytic in-gel digestion mass spectrometry revealed that ECSIT_CTER_ was multiphosphorylated (**Fig. S10, Table S2**). We were able to identify nine high-confidence Threonine/Serine phosphorylation sites, including a kinase recognition sequence TSS motif ^23^ (res. 401-403) (**Fig. 4C, Fig. S10, Table S2**).

**Figure 4.**
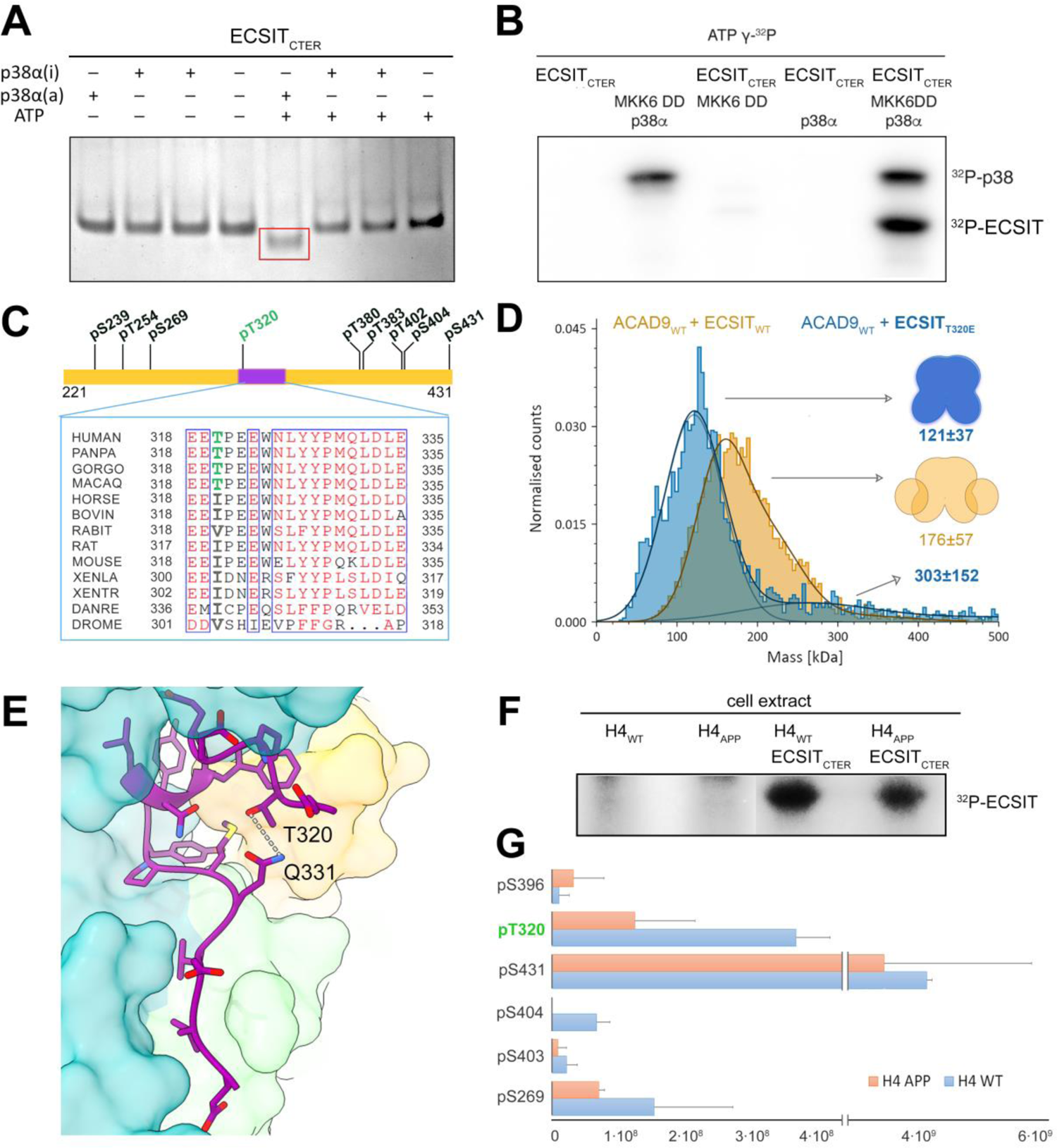
Phosphorylation of ECSIT may regulate the binding affinity for ACAD9. **(A)** ECSIT treated with MKK6-activated p38α MAP kinase reveals a different mobility shift compared to the non-treated protein on a native gel. **(B)** Radioblotting assays of active P38α MAP kinase and ECSIT p38α-treated sample corroborate the ECSIT phosphorylation forms. **(C)** *Top*, diagram summarising the phosphorylation sites of ECSIT_CTER_ identified by mass spectrometry. The sequence observed in the cryo-EM map is highlighted in purple and the potential Thr320 phosphorylation site in green. Multiple sequence alignment of this region between different species reveals a high conservation. However, Thr320 seems to have appeared in higher primates as a replacement of a hydrophobic residue, suggesting that this substitution might have appeared as an evolutionary shift in function. **(D)** Mass photometry reveals that the ECSIT_T320E_ phosphomimetic mutant affects the complex formation with ACAD9, resulting in a large particle with the major species being an ACAD9 homodimer with unbound ECSIT. **(E)** Structural details showing the interaction of Thr320 with Gln331 in the ECSIT_CTER_ segment. A phosphate group in Thr320 would generate a steric clash that would result in a no longer compatible conformation for binding to ACAD9. **(F)** Cell extracts from neuronal cells H4, both wild-type (WT) and amyloidogenic (APP), incubated with ECSIT_CTER_ confirmed the phosphorylation of ECSIT_CTER_ *ex cellulo* by a kinase pool. **(G)** Quantitative mapping of the ECSIT_CTER_ phosphopeptides indicate a decreased phosphorylation level under amyloidogenic conditions, with Thr320 (in green) displaying the highest differential level.

Excitingly, one of the identified potential phosphorylation sites was Thr320, which resides within the ECSIT sequence visible in our ACAD9-ECSIT_CTER_ structure (**Fig. 4C, E**). Thr320 forms hydrogen bonds with Gln331, stabilising the helical sequence that enters into the ACAD9 binding cavity. We therefore constructed an ECSIT_CTER_ phosphomimetic mutant where Thr320 was replaced by a Glutamate (T320E). Remarkably, this mutant exhibited a significantly reduced binding affinity for ACAD9 (**Fig. 4D**), suggesting that the phosphorylation of Thr320 may negatively regulate ACAD9 binding.

We next sought to examine the potential ECSIT_CTER_ phosphorylation sites *in vivo*. To this end, we treated ECSIT_CTER_ with an endogenous kinase pool from human neuronal cells (**Fig. 4F**). We observed strong ^32^P-radiolabelling of ECSIT_CTER_, providing evidence that ECSIT undergoes phosphorylation *ex cellulo*. Subsequent analyses by proteolytic in-gel digestion mass spectrometry confirmed that ECSIT_CTER_ was multiphosphorylated, and identified six high-confidence Threonine/Serine sites, including five already detected *in vitro*: Ser269, Thr320, the TSS motif (401 to 403) and the C-terminal residue Ser431 (**Fig. 4C, F-G, Table S3**).

### ECSIT_CTER_ phosphorylation is affected by Aβ soluble oligomers

In parallel, we performed a quantitative phosphoproteomic study upon exposure of ECSIT_CTER_ to soluble Aβ oligomers (**Fig. 4F-G**). Intra-neuronal Aβ can translocate directly from the endoplasmic reticulum into mitochondria, where it may affect mitochondrial respiration^24^. Thus, we sought to investigate whether mitochondrial Aβ alters the phosphorylation levels of ECSIT. Overexpression of the amyloid precursor protein (APP) carrying the AD-related Swedish mutation (KM670/671NL) in neuroglioma cells enables the investigation of the toxic Aβ_1-42_ soluble oligomeric form as may occur during AD^25, 26^ (**Fig. S11A**). Interestingly, while we obtained similar levels of immunopurified CI from both cell types (**Fig. S11B, *inset***), the amyloidogenic cells exhibited a significantly enhanced NADH-dehydrogenase activity (**Fig. S11B**), raising the possibility that soluble Aβ_1-42_ oligomers lead to aberrant CI hyperactivity and could be a primary source of oxidative stress. Finally, we investigated whether the Aβ_1-42_ soluble oligomers affect ECSIT_CTER_ phosphorylation and found that the degree of phosphorylation of the six phosphorylation sites differed greatly between wild-type and amyloidogenic neuronal cells, with a clear tendency to decrease upon exposure to amyloids (**Fig. 4G, Table S3**).

## DISCUSSION

The mitochondrial Complex I assembly complex (MCIA) is required for the biogenesis of Complex I and is therefore crucial for the activation of the OXPHOS system^5^. FAO and OXPHOS are key pathways involved in cellular energetics^27^. Despite their functional relationship, evidence for a physical interaction between the two pathways is sparse^28^. Understanding how FAO and OXPHOS proteins interact and how defects in these two metabolic pathways contribute to mitochondrial disease pathogenesis is thus of critical importance for the development of new tailored therapeutic strategies.

Here, we provide high-resolution structural insights into the interface and assembly of the MCIA subcomplex ACAD9-ECSIT, with one molecule of ECSIT bound to each ACAD9 protomer at the crossing between the dehydrogenase and vestigial domains. ECSIT induces the flipping of the ACAD9 β1-β2 loop acting as gatekeeper by ∼10 Å such as to allow the binding of an ECSIT segment (res. 320-334) and the release of the FAD cofactor from ACAD9 (**Fig. 3**). Once formed, the complex is very stable (**Fig. S1A-D**). Based on the present data and our previously published low resolution information^11^, we propose this stability may arise from ECSITs ability to envelop ACAD9, made possible by long stretches of flexible loops (res. 306-320 and 335-356, **Fig. S4A-C**).

Furthermore, the AF2-predicted ACAD9-ECSIT_CTER_ model supports our previous suggestions that ECSIT entwines around the vestigial domain of ACAD9 (**Fig. S4A, C**)^11^, in contrast to an alternative SAXS model placing the ECSIT binding interface at the N-terminus of ACAD9^12^. The fact that there is an overall low confidence in the position of the globular core of C-terminal ECSIT with respect to ACAD9 (**Fig. S4B**) not only supports our experimental data but may also explain why the folded region of ECSIT is not visible in the ACAD9-ECSIT_CTER_ reconstruction: this region is probably highly mobile due to the flexibility of residues 306-320 and 335-356. Worthy of note, because AF2 is unable to predict conformational changes that occur during protein-protein interactions^29^, while the AF2 model corroborates the experimentally determined ACAD9-ECSIT_CTER_ interface, it does not predict the displacement of the gatekeeper loop.

The experiments with the ECSIT-mimicking short peptide identified from the cryo-EM map show that this stretch of residues is key to the ejection of the FAD cofactor from ACAD9 (**Fig. 1D**). Intriguingly, despite the high similarity between the ACAD9 and VLCAD structures, only ACAD9 holds the unique ability to bind ECSIT and participate in CI assembly^6^. In order to investigate key differential residues, we first examined the divergent residues on the gatekeeper loop. However, mutant analyses reveal that, even if the sequence of this loop has an impact in the acyl-CoA activity of the enzyme, the exact nature of the amino acids is not directly responsible for the formation of the complex with ECSIT (**Fig. S7B-D**). We then examined the FAD binding site in both protein structures^15^ (**Fig. S8A-B**) to check whether the FAD binding stability is related to the ECSIT binding capacity. We observed that in fact, ACAD9 has a larger FAD binding pocket, leading to a reduced number of bonding interactions (**Fig. S8A-B**) that are typically longer and therefore, weaker (**Fig. S8A-D**). A less tightly bound cofactor is less stable and thus the feasibility of ejection should be higher than if the FAD was tightly bound. Very interestingly, through the identification of VLCAD residues in the FAD binding, we have reversed the ACAD9 behaviour in the counterpart mutant, achieving reduction both in ECSIT binding and in FAD ejection (**Fig. S8C-G**). In fact, the counterpart mutation in VLCAD also induces a reversed behaviour, facilitating ECSIT binding to VLCAD (**Fig. S8B, D, F-G)**. These findings suggest that changes in hydrophobicity/π-bonding capabilities in the FAD binding pocket strongly affect the FAD stability and indirectly, the binding to ECSIT as well (**Fig. S8B-G**). Furthermore, re-examination of a VLCAD structure at higher resolution and in absence of fatty acid substrate revealed a unique quaternary structure of VLCAD homodimers stabilised by an extended conformation of the ∼35 residue stretch at the base of VLCAD (486-521, **Fig. S9A**) in contrast to the equivalent residues in ACAD9 (**Fig. S9B-C**)^18^. Notably, this region is also where both proteins show the highest sequence divergence, which could further account for dissimilar conformational abilities.

Remarkably, this extended loop occupies the equivalent ‘ECSIT binding site’ (**Fig. S9D-E**), suggesting that a reason why VLCAD is unable to bind ECSIT may be the obstruction of this binding site by VLCAD itself. Taken together, subtle differences in sequence are thus likely to play a critical role in impacting ACAD9-ACAD9 and VLCAD-ECSIT interactions, possibly accounting for the differences between VLCAD and ACAD9 in CI assembly participation, as previously suggested^6^.

Our finding that ECSIT undergoes phosphorylation raises the possibility that a post-translational modification may provide an additional layer of regulation of MCIA complex assembly (**Fig. 4**). Although further investigation is required to determine the identity of the ECSIT kinases and the mechanism by which these might regulate MCIA complex formation, the identification of five high-confidence Threonine/Serine sites in ECSIT_CTER_ suggests a potential role for phosphorylation in modulating the ACAD9-ECSIT interaction. In particular, our analyses of the phosphomimetic mutant T320E suggest that phosphorylation at Thr320, either solely or in combination with other phosphorylation sites, could conceivably induce conformational changes that would disrupt the fitting of the ECSIT segment in the ACAD9 cavity (**Fig. 4E**). Notably, analysis of the segment sequence across species reveals a high amino acid conservation except for Thr320, which seems to have evolved in higher primates from an Isoleucine (**Fig. 4C**). Along this line, it is interesting to higlight that Threonine is an intermediate in Isoleucine biosynthetic pathway, which supports the hypothesis that rapid evolution of phosphorylation sites could provide a way to fit the environment by rewiring the regulation of signal pathways. Furthermore, previous studies have found evidence that the functional potential of phosphorylation sites are increased with their evolutionary age, especially for Serine and Threonine, in disordered regions^30^.

Finally, our observations of decreased ECSIT_CTER_ phoshorylation in response to Aβ oligomers, coupled to a significantly enhanced NADH-dehydrogenase CI activity, further suggests that ECSIT plays a role as a molecular link between mitochondrial bioenergetics and early amyloid pathology (**Fig. 4G**, **Fig. S11**). Although further investigation is needed to determine how amyloids contribute to the functional activities of ECSIT, soluble amyloids would somehow favor ECSIT dephosphorylation, apparently required for the initial recognition of MCIA partners. Under early amyloidogenic conditions, a stabilised MCIA complex would in turn contribute to a stabilised CI and thus, enhance its activity. Progressively, CI over-activity would lead to an NADH/NAD+ redox imbalance, generating oxidative stress and exacerbating the accumulation of Aβ oligomers in a vicious cycle, which ultimately may result in the inhibition of CI activity, in agreement with previous studies^2, 20^.

## CONCLUSION

Taken together, our study provides insights into the molecular mechanisms of ACAD9-ECSIT assembly and its functional implications in FAO and OXPHOS pathways, a process that may be in turn regulated by the phosphorylation of ECSIT (**Fig. 5**). Here, in the absence of ECSIT, ACAD9 acts as an acyl-CoA dehydrogenase enzyme in the first step of the FAO pathway. The recognition of aromatic residues on the ECSIT 320-334 residue segment by the ACAD9 gatekeeper loop would induce its opening, enabling the ejection of the FAD cofactor and the binding of ECSIT. This dual mechanism seems to be unique to ACAD9 by virtue of the flexible nature of its FAD binding site and the compactness of the 35 amino acid stretch at the base of the molecule. Our findings that ECSIT can be phosphorylated and that this has a negative effect on its affinity for ACAD9 imply that an ECSIT (de-)phosphorylation would be the trigger of ECSIT binding to ACAD9 and hence, a regulatory switch of the MCIA complex assembly. The observations that Aβ soluble oligomers increase the activity of CI but also decrease ECSIT phosphorylation levels provide further evidence that ECSIT might be involved in AD pathogenesis^10^. By compromising both the assembly of CI and the regulation of FAO, ECSIT could act as a reprogrammer of OXPHOS and FAO pathways that may lead to altered metabolism in brain mitochondria and ultimately neurodegeneration^11^.

**Figure 5.**
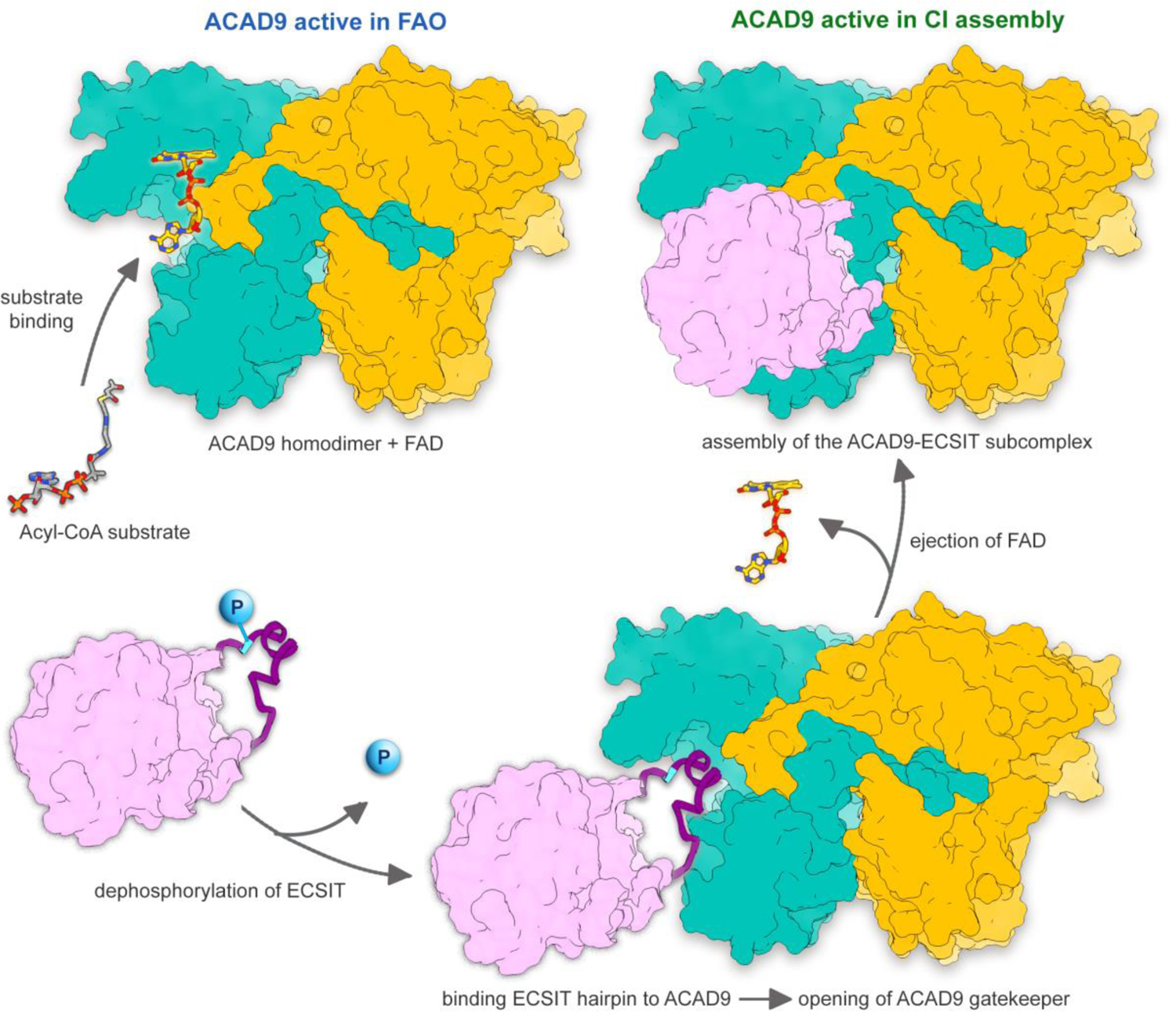
Proposed model of the mechanism of ACAD9-ECSIT assembly and its functional implications in FAO and OXPHOS pathways. In absence of ECSIT, ACAD9 acts as an acyl-CoA dehydrogenase enzyme in the first step of the FAO pathway. Dehydrogenation of the acyl-CoA substrate is concomitant with the reduction of the FAD prosthetic group into FADH_2_. Upon recognition of ACAD9 by ECSIT, the gatekeeper loop in ACAD9 flips upwards, allowing the concomitant binding of ECSIT and ejection of FAD, shutting down the dehydrogenase activity of ACAD9 and becoming committed to CI assembly. ECSIT dephosphorylation would enable the binding to ACAD9 and hence, act as a potential trigger to favour the assembly of the MCIA complex.

## METHODS

### DNA plasmids

The DNA plasmids were constructed following the same protocol as in ^11^. Mutant plasmids were generated using the QuikChange Lightning Site-Directed Mutagenesis Kit (Agilent). ACADVL was a gift from Nicola Burgess-Brown (Addgene plasmid #38838; http://n2t.net/addgene:38838). ACAD9-related constructs were cloned into pET21d while VLCAD-related constructs were cloned into pNIC28-BSA.

### Recombinant protein expression and purification

ECSIT_CTER_ (residues 221-431) and all ECSIT mutants (ECSIT_W324A_, ECSIT_Y327F_, ECSIT_Y327A_, ECSIT_Y328F_, ECSIT_Y328A_, ECSIT_T320E_) were expressed in *E. coli* BL21 Star (DE3) cells (ThermoFisher Scientific) in LB medium at 25 °C and subsequently induced with 250 μM isopropyl-B-galactopyranoside (IPTG). ACAD9_WT_ (res. 38-621) and ACAD9 mutants (ACAD9_S191A_, ACAD9_A184S_, ACAD9_SR194TS_, ACAD9_A184S_/_SR194TS_, ACAD9_VLCAD_) were expressed in *E. coli* C43 (DE3) cells (Lucigen) while VLCAD_WT_ (res. 72-655) and VLCAD_ACAD9_ in *E. coli* Rosetta 2 (DE3) cells (Merck), and cultured in Terrific Broth medium at 37 °C and induced with IPTG at a final concentration of 500 μM. All cells were harvested 18 hours after induction. Bacterial pellets were resuspended in a lysis buffer (200 mM potassium phosphate, 300 mM or 250 mM NaCl, 0.25 mM EDTA, 1 mM DTT, 0.2% Tween-20, pH 7.5) supplemented with protease inhibitor cocktails (Merck) and DNase I (Merck). Bacterial lysis was performed by sonication (QSonica) followed by removal of cell debris by centrifugation for 25 minutes, 45 000g (Avanti J-20 XP centrifuge, Beckman Coulter) at 4 °C. Protein purification was conducted using a combination of affinity and size-exclusion chromatography (SEC) performed using Äkta Purification systems (GE Healthcare). Lysate containing the protein of interest was loaded onto a 5 mL HisTrap column (GE Healthcare) equilibrated in 200 mM potassium phosphate, 300 mM or 250 mM NaCl, 5 mM BME, pH 7.5 and eluted with a linear imidazole gradient reaching a final concentration of 500 mM imidazole in 200 mM potassium phosphate, 300 mM or 250 mM NaCl, 5 BME, pH 7.5. Protein-containing fractions were identified using sodium dodecyl sulfate-polyacrylamide gel electrophoresis (SDS-PAGE) and selected for further purification. VLCAD (and mutant) and ECSIT_CTER_ (and mutants) were purified on Superdex 200 10/300 GL SEC column (GE Healthcare), equilibrated in 25 mM HEPES, 300 mM or 250 mM NaCl, 1 mM TCEP, pH 7.5. ACAD9_WT_ (and mutants) were dialysed in SEC buffer. Sample purity was assessed using SDS-PAGE and concentrated using 30 kDa and 10kDa cut-off Amicon Ultra filters for ACAD9/VLCAD and ECSIT, respectively. Single proteins were stored at −80 °C until use.

The MCIA subcomplex ACAD9-ECSIT (all variants) was assembled by warming an aliquot of ACAD9 to 32°C and slowly adding ECSIT to a final molar ratio of 1:1.25. After ∼5 minutes the protein mixture was plunged into ice and left for at least 15 minutes before use. Protein complex samples were used in analyses immediately after formation. Constitutively active MKK6DD (S207D/T211D mutant) and WT p38α were produced as described in ^31^.

### Isothermal Titration Calorimetry (ITC)

Measurements were done on a Micro PEAQ-ITC (Malvern Panalytical) at 25 °C. ACAD9_WT_, ECSIT_CTER_, ECSIT_T320E_ were dialyzed overnight against 50mM TRIS pH 7.5, 250mM NaCl. For binding studies, ACAD9_WT_ at 49 µM was introduced into the sample cell and titrated with either ECSIT_CTER_ or ECSIT_T320E_ respectively at 300 µM over a period of 50 min. Data analysis was carried out using the MicroCal PEAQ-ITC Analysis Software (Malvern Panalytical). The experiment was performed in triplicate.

### Mass Photometry data acquisition and analysis

Mass Photometry experiments were carried out using a Refeyn OneMP Mass photometer system (Refeyn Ltd, Oxford, UK). All data were acquired with a 3*10 µm (at 1 kHz) size field of view. AcquireMP and DiscoverMP software packages were used to record movies and analyse data respectively using standard settings. Microscope coverslips (high precision glass coverslips, Marienfeld) were cleaned following the Refeyn Ltd Individual rinsing procedure. Reusable self-adhesive silicone culture wells (Grace Bio-Labs reusable CultureWell^TM^ gaskets) were used to keep the sample droplet shape. Contrast-to-mass calibration was carried out using a mixture of native proteins (NativeMark Unstained Protein Standard, Invitrogen) with molecular weight (MW) of 66, 146, 480 and 1048 kD. Immediately prior to the measurements, protein stocks were diluted directly in analysis buffer (25 mM HEPES pH 7.5, 250 mM NaCl), or in 4.8 µM of ECSIT solution in case of complex formation, to reach a concentration of 20-50 nM for ACAD9. To this end, 1 µL of protein solution was added into 19 µl of analysis buffer, to reach a final drop volume of 20 µL.

### Size Exclusion Chromatography coupled to Multi-Angle Laser Light Scattering (SEC-MALLS)

The molecular mass of ECSIT_CTER_ and ACAD9_WT_ in solution was determined by SEC coupled to multi-angle laser light scattering (SEC-MALLS) using a Superdex 200 10/300 GL column equilibrated in 25 mM HEPES, 250 mM NaCl, pH 7.8 buffer. The measurements were performed at 20 °C using 50 μL of proteins at 5 mg/mL with a flow rate of 0.5 mL/min and eluted proteins were monitored using a DAWN-EOS detector with a laser emitting at 690 nm for online MALLS measurement (Wyatt Technology) and with a RI2000 detector for online refractive index measurements (SchambeckSFD). Molecular mass calculations were performed with the ASTRA software using a refractive index increment (dn/dc) of 0.185 mL/g. Data were visualised using OriginPro 9.0 (OriginLab) software.

### Dynamic Light Scattering (DLS)

DLS measurement were performed at 278K on a Zetasizer Nano S (Malvern Panalytical), using 20 µl of protein sample at 1-2 mg/ml. A Micro Cell quartz cuvette was incubated at 4 °C until a stable temperature was reached. 13 replicas (comprising 7 measurements) were done for each sample. Data was processed using Zetasizer Software 8.01.4906.

### Acyl-CoA dehydrogenase (ACAD) activity assay

The assay was performed as previously described ^11^.

### Cryo-EM sample preparation

ACAD9_WT_-ECSIT_CTER_ was purified through gel filtration as described above and stored in a buffer containing 25mM HEPES, 300 mM NaCl and 1 mM TCEP, pH 7.5. The ACAD9_WT_ and ACAD9_S191A_ samples were dialysed into the same buffer and stored without gel filtration. For cryo-EM grid preparation, the samples were diluted in the same buffer to 0.17 mg/mL, 0.03 mg/L and 0.02 mg/mL respectively, applied to a glow-discharged R 1.2/1.3 300 mesh UltrAuFoil gold grid (Quantifoil Micro Tools GmbH) and plunge-frozen in liquid ethane using a Vitrobot Mark IV (Thermo Scientific) operated at 8°C at 100 % humidity.

### Cryo-EM data collection

Grids were pre-screened on a Glacios microscope (Thermo Scientific) of the EM platform of the Institute of Structural Biology (IBS), Grenoble, France. Datasets were collected at the European Synchrotron Radiation Facility CM01, on a Titan Krios microscope (Thermo Scientific) operating at 300 kV with a Quantum energy filter (slit width 20 eV) ^32^. The ACAD9_WT_-ECSIT_CTER_ and ACAD9_WT_ data sets were recorded at zero degrees stage tilt on a K2 summit direct electron detector (Gatan) running in counting mode, at a magnification of 165,000x, corresponding to a pixel size of 0.827 Å/pixel at the specimen level. The ACAD9_S191A_ data set was recorded at a 35° stage tilt, on a K3 direct electron detector (Gatan) operating in super-resolution mode, at a magnification of 105,000x, with a sampling pixel size of 0.42 Å/pixel, and was binned two-fold for data processing. For ACAD9_WT_-ECSIT_CTER_, a total of 8,001 movies of 40 frames were acquired with a dose rate of 9.5 e-/Å^2^/s and a total exposure time of 3 s, corresponding to a total dose of 41.7 e-/Å^2^. For ACAD9_WT_, a total of 7,731 40 frames-movies were acquired with a dose rate of 6.7 e-/Å^2^/s and a total exposure time of 6 s, corresponding to a total dose of 40.2 e-/Å^2^. For ACAD9_S191A_, a total of 2,641 40 frames-movies were acquired with a dose rate of 14.7 e-/Å^2^/s and a total exposure time of 3 s, corresponding to a total dose of 62.5 e-/Å^2^.

### Cryo-EM data processing

All image analysis was conducted using the cryoSPARC software^33^. Imported movies were motion-corrected, dose weighted, and, in the case of the super-resolution ACAD9_S191A_ data set, Fourier cropped (2x). All frames of the ACAD9_WT_-ECSIT_CTER_ and ACAD9_WT_ movies were used, whereas the first 5 frames of the ACAD9_S191A_ movies had to be discarded due to a stage drift during data acquisition at 35° tilt. Initial CTF estimation was performed on the aligned and dose-weighted summed frames. Micrographs were then manually screened, resulting in 7,789 micrographs of ACAD9_WT_-ECSIT_CTER_, 7,598 micrographs of ACAD9_WT_ and 2,608 micrographs of ACAD9_S191A_ selected for further processing.

The ACAD9_WT_-ECSIT_CTER_ data set was the first to be acquired and processed. The first round of particle picking was performed automatically using the blob picker, resulting in ∼2,200,000 picked particles. Particles were then extracted with a box size of 256 × 256 pixels and subjected to 2D classification, which immediately revealed very strongly predominant C2-symmetric ‘top view’ orientation, similar to the situation encountered in our earlier work with a different ECSIT construct ^11^. ∼1,600,000 particles corresponding to the best classes showing clear secondary structural features were selected for the generation of the first *ab initio* 3D volume which was then used as a reference for homogeneous refinement with applied C2 symmetry. Although the resulting 3D reconstruction had a nominal resolution of 3.07 Å, it was clearly anisotropic due to the strong preferential ‘top view’ orientation and the under-representation of the side and tilted views. Thus, projections of this reconstruction according to the directions of rare views were selected as templates for the next round of particle picking, leading to ∼2,500,000 picked particles. These were extracted and subjected to 2D classification resulting in a more diverse set of classes. Seven classes (∼115,000 particles, **Fig. S2A**) representing the most different views of the molecule were used for a homogenous 3D refinement with the previous reconstruction as a reference and a C2 symmetry imposed, which yielded a map at 3.23 Å. This improved but still visually somewhat elongated map was used to generate templates corresponding to different subsets of the rarest views for several further picking and 2D classification jobs performed in parallel. Particles corresponding to the rare views were combined, duplicate particles resulting from different picking jobs removed with a dedicated cryoSPARC routine, and classes representing the most different orientations carefully selected for a 3D refinement, resulting in a new 3D volume and leading to generation of new rare view templates. This procedure was iterated until no further improvement of the reconstructed 3D volume was detected. This 3D volume was then used as a reference for homogeneous 3D refinement of the corresponding 279,483 particles, followed by a non-uniform 3D refinement. This last refinement step resulted in a final 3D map with an estimated resolution of 3.07 Å according to the Fourier Shell Correlation (FSC = 0.143), which was then post-processed using a soft mask and sharpened with a B-factor of −171 Å^2^ for visualisation and model building (**Fig. 1B, S2B-D**).

The second acquired and processed data set, ACAD9_WT_, suffered from an apparently even more severe preferential orientation. Therefore, after numerous trials with the blob picker and the templates obtained from the resulting 2D class averages, we resolved to use the rare views of the final ACAD9_WT_-ECSIT_CTER_ map filtered to 30 Å resolution for template picking. A total ∼1,120,000 particles were picked in several parallel jobs and duplicates removed. The particles were extracted with a box size of 256 x 256 pixels and subjected to 2D classification. Homogenous 3D refinement with the low pass filtered ACAD9_WT_-ECSIT_CTER_ map as a reference and imposing the C2 symmetry was then performed on several different subsets of classes while carefully selecting rare views of reasonable quality. The best, although anisotropic, reconstruction that could be obtained by this approach included only 31,209 particles and had a nominal resolution of 3.91 Å. These particles were then subjected to a 3D classification resulting in two classes containing 17,922 and 13,287 particles, and corresponding 3D maps with estimated average resolutions of 4.71 Å and 5.98 Å, respectively. Visual assessment of these maps showed that the lower resolution map containing less particles was actually of higher quality and had a more isotropic resolution (**Fig. S5A-D**). Homogeneous 3D refinement of this map showed a further improvement in its quality, leading to a final 3D reconstruction of ACAD9_WT_ with an estimated average resolution of 4.5 Å (FSC = 0.143). This map was post-processed using a soft mask and sharpened with a B-factor of −180 Å^2^.

Considering the difficulties encountered due to a very strongly preferred orientation of ACAD9_WT_, we opted for a recording of the ACAD9_S191A_ data set at a 35 degrees stage tilt in order to increase the proportion of the rare views. As for ACAD9_WT_, the rare views of the low pass filtered ACAD9_WT_-ECSIT_CTER_ map were used for template picking. A total ∼286,000 particles were picked, extracted with a box size of 256 x 256 pixels and subjected to 2D classification. The best classes containing a total of ∼149,000 particles were selected (**Fig. S5E**) and subjected to a homogenous 3D refinement, using the low pass filtered ACAD9_WT_-ECSIT_CTER_ map as a reference and imposing the C2 symmetry. The obtained volume had a nominal resolution of 4.02 Å but, similarly to what we observed for the first map of ACAD9_WT_-ECSIT_CTER_, clearly anisotropic. Addition of ∼14,000 more particles corresponding to other classes resembling the rarer views and a subsequent 3D refinement resulted in a map of 3.92 Å resolution. A non-uniform 3D refinement of these ∼163,000 particles gave a map with an estimated resolution of 3.63 Å (FSC = 0.143).

A dip in the FSC curve at ∼ 6.5 Å was however still indicative of a remaining anisotropy (**Fig. S5F**). In addition, a visual inspection of the map showed that some regions, in particular the β-sheets were not resolvable. However, side chain information could be identified in most α-helices. Further data processing with additional particles and changing parameters showed no improvement in the map quality (**Fig. S5G-H**). Thus, this latest 3D reconstruction of ACAD9 _S191A_ was manually sharpened with an input B factor of −180 Å^2^ to allow better visualisation of the higher resolution features such as side chains and the orientation of the FAD cofactor.

### Model building and refinement

To build a model of ACAD9_WT_-ECSIT_CTER_, a homology model of ACAD9 dimer based on the human very long-chain Acyl-CoA dehydrogenase (VLCAD, PDB:3b96), was used as the initial structure, as described in ^6^. AF2 models were not available at the time. In this model, each ACAD9 protomer contains residues 38-621 and binds one FAD molecule (a total of two FADs in the ACAD9 dimer). The homology model was rigid-body fitted into the 3D reconstruction of ACAD9_WT_-ECSIT_CTER_ using ChimeraX ^34^. A first round of real space refinement was conducted enabling rigid body, global minimization, local grid search and atomic displacement parameter (ADP) refinement parameters; rotamer Ramachandran, non-crystallographic symmetry (NCS) and reference model (VLCAD-based homology model) restraints were also imposed. The output model was then manually corrected in Coot ^35^ followed by iterative real space refinements with PHENIX ^36^, using the same settings but removing the rigid body refinement parameter.

Regions of the ACAD9 dimer model with no clearly corresponding cryo-EM density (454-486, FAD) were removed from the final model. Upon model building, additional density that could not be attributed to ACAD9 was observed in a pocket adjacent to the ACAD9 dehydrogenase domain on each protomer. We tentatively attributed this density to ECSIT and manually inserted short polyalanine chains into each density using Coot ^35^. Each chain was seen to adopt a short α-helix composed of 8-10 residues, sandwiched between two loops. A total of 15 Alanine residues could be fitted into the visible density. Due to the high resolution of the cryo-EM map in this region, it was possible to identify characteristics of several aromatic side chains, in particular a Phenylalanine and two neighbouring Tyrosine residues, indicating a sequence WxYY where X represented an unidentified residue. Comparison with the sequence of the ECSIT construct enabled us to detect a short amino acid sequence, totalling 15 amino acids corresponding to residues 320-334, with aromatic side chain positions matching those observed in the ACAD9_WT_-ECSIT_CTER_ cryo-EM map. The sequence was then mutated in Coot ^35^ in agreement with the identified ECSIT sequence. The final ACAD9_WT_-ECSIT_CTER_ model was validated using the comprehensive validation tool in PHENIX ^36^.

Considering that the final cryo-EM map of ACAD9_WT_ was at 4.5 Å resolution and showed high levels of anisotropy, no attempts were made to build an atomic model of ACAD9_WT_ in the absence of ECSIT. Thus, a simple rigid body fit of the ACAD9_WT_-ECSIT_CTER_ model and the AF2 ACAD9 model (res. 38-621) was performed in ChimeraX ^34^. Model building into the ACAD9_S191A_ cryo-EM map was done using both the VLCAD-based ACAD9 homology model and the final ACAD9_WT_-ECSIT_CTER_ models in parallel. Rigid body fitting in ChimeraX revealed that the ACAD9_WT_-ECSIT_CTER_ model fitted better into the ACAD9_S191A_ cryo-EM map than the VLCAD-based ACAD9 homology model. Therefore, the ACAD9_WT_-ECSIT_CTER_ model was used for further structure refinement. As demonstrated in our earlier work ^11^, the FAD is not present in the ACAD9_WT_-ECSIT_CTER_ cryo-EM map. Therefore, in the resulting atomic model, the coordinates of the FAD molecules were taken from the aligned PDB of the homology model. Similarly to ACAD9_WT_-ECSIT_CTER_, the ACAD9_S191A_ cryo-EM map does not display density for the flexible helices located next to the vestigial domain of ACAD9 (residues 454-485). As a result, these were removed from the model. Real space refinement of secondary structural features as well as side chains was iteratively conducted in Coot followed by rigid body refinement using PHENIX ^36^. The final ACAD9_S191A_ model was also validated using the comprehensive validation tool in PHENIX ^36^. A summary of refinement and model validation statistics can be found in **Table S1**.

### AlphaFold modelling

Structures of ACAD9 (human, residues 1-621, monomer) and ECSIT (human, residues 1-431) were available in the AlphaFold Protein Structure Database (AF2)^16, 29^. The PDB files were downloaded and compared to the structures produced from experimental data. For the structural comparison of ECSIT model (residues 320-334) built based on the cryo-EM map, the AF2 ECSIT model was cropped to match the sequence of our construct without the addition of His-Tags. The structures were aligned and RMSD values generated using ChimeraX^34^.

AlphaFold modelling of the ACAD9-ECSIT subcomplex was conducted using ColabFold prediction tools^37^. The protein sequences for ACAD9 (human) and ECSIT (human) were taken from UniPROT^38^ and modified to match the constructs used in our experiments: ACAD9 contained residues 38-621 and ECSIT contained residues 221-431. Structure prediction of the ACAD9-ECSIT subcomplex was performed using two ACAD9 sequences and two ECSIT sequences, based on information obtained from our experimental results.

We took the best predicted model from the five proposed models for our analysis. The Predicted Aligned Error (PAE) plot generated alongside the docked model showed low values (< 15), representing well defined relative positions and orientations (**Fig. S4**), for the majority of the residues in the ACAD9 dimer, the exception being for residues 455-486 which have high PAE values (∼30). Overall PAE values for ECSIT are high, however lower values are shown for the residues corresponding to those modelled in the ACAD9_WT_-ECSIT_CTER_ model, where the PAE value is low. Additionally, alignment of the ECSIT residues built based on the cryo-EM map with the calculated model shows a very high agreement (RMSD of 1.045 Å) (**Fig. S4D**), providing further support to our experimental results. **Fig S4** describing the AF results was generated using ChimeraX^34^.

### Mammalian cell culturing

Human neuroglioma H4 cells, both wild-type (WT) and stably transfected with human amyloid precursor protein (APP) carrying the AD-related Swedish mutation (KM670/671NL)^39^, were kindly provided by Dr. Patrick Aloy, Institute for Research in Biomedicine, Barcelona. Cells were cultured in Dulbecco’s Modified Eagle Medium (DMEM)-low glucose and supplemented with GlutaMAX, pyruvate (Thermofisher), 10%(v/v) fetal bovine serum (FBS, GIBCO) and 100 U/mL penicillin–streptomycin (Invitrogen) at 37 °C and 5% CO2 in a humidified incubator.

For mitochondrial assays, cells were trypsinized at 80% confluence after 72-hour incubation on T225 flasks (10^6^ cells / flask). For ECSIT_CTER_ phosphorylation assays, cells were cultured in galactose-free media and trypsinized at 80 % confluence after 4-hour incubation on T225 flasks (10^6^ cells / flask).

### Mitochondria isolation, CI immunopurification, CI activity assay and mitochondrial Aβ_1-42_ detection

About 25 x 10^6^ cells were resuspended in homogenization buffer (10 mM Tris-HCl pH 6.7, 10 mM KCl, 0.15 mM MgCl2, 1 mM PMSF, 1 mM DTT) and then transferred to a glass homogenizer and incubated for 10 min on ice. Cells were lysed using a tight-fitting pestle for approximately 2 min. The homogenized cellular extract was then gently mixed with 2 M Sucrose and centrifuged three times at 1,200 × g for 5 min to obtain the supernatant. Mitochondria were isolated by differential centrifugation steps according to published protocols^40^. Mitochondria were pelleted by centrifugation at 7,000 × g for 10 min and centrifuged twice in wash buffer (250 mM sucrose, 10 mM TRIS HCl, pH 6.7, 0.15 mM MgCl2, 1 mM PMSF and 1 mM DTT) at 9,500 × g for 5 minutes. Complex I (CI) was immunopurified from H4 cell mitochondria using a commercial kit (Abcam ab109721) according to the manufacturer instructions. All steps were carried out at 4 °C. Briefly, mitochondria were solubilised with 1% n-dodecyl β-D-maltoside (DDM), centrifuged to remove insoluble material and incubated with the affinity beads overnight. The beads were washed twice before CI was eluted in buffer containing 200 mM glycine (pH 2.5) and 0.5% DDM. The pH was neutralised by addition of Tris base. Solubilised mitochondria were resuspended in PBS buffer containing protease inhibitors (cOmpleteTM, Sigma) and concentration adjusted to 5.5 mg/mL using Bradford assay (BioRad). 1 % DDM was then added, the preparation incubated on ice for 30 min and centrifuged at 16,000 x g for 10 min. Samples were added to the microplate wells precoated with a specific CI capture antibody. CI activity was analysed by measuring the absorbance at OD450 nm in a kinetic mode at room temperature for up to 16 hours in a microplate reader Quantamaster QM4CW (Horiba). The human Amyloid Beta (residues 1-42, Aβ_1-42_) content in isolated mitochondria from WT and APP cells was measured by enzyme-linked immunosorbent assay (ELISA) according to the manufacturer instructions (ThermoFisher khb3544).

Every assay was carried out with three independent experiments and presented as a mean average with the standard deviation. Statistically significant differences were determined by one-way ANOVA followed by Tukey-Kramer post-test to identify pair wise differences. Differences were considered significant at P < 0.01(**), P < 0.0001 (****). Statistical analyses were carried out using GraphPad Prism version 8 (GraphPad Software).

### Phosphorylation assays *in vitro*

P38α MAP kinase was activated with the active (DD) MKK6 kinase form. Protein samples were prepared on ice (MKK6DD:p38α:ECSIT) in 10 µl phosphorylation reaction buffer. For radiolabelling experiments, radiolabelled nucleotide ATP γ-^32^P (Hartmann Analytic GmbH)) was diluted 1:10 into a 1 mM cold ATP solution. The phosphorylation reaction was started by adding 1 μl of nucleotide to each sample and incubated for 20 min at 30°C. The reaction was stopped by adding 4 μl of SDS sample buffer (0.4% bromophenol blue, 0.4 M DTT, 0.2 M Tris pH 6.8, 8% SDS, 40% glycerol) and boiling for 5 min at 95 ℃. Samples were centrifuged and loaded on Pre-cast 4-20% gradient Tris-Glycine gels (ThermoFisher Scientific), run in Tris-Glycine running buffer (2.5 mM Tris Base, 19.2mM glycine pH 8.3, 1% SDS) at 220 V for 40 minutes. The gels were exposed to a storage phosphor screens (GE Healthcare) overnight and imaged using a Typhoon scanner (GE Healthcare). Gels were then stained with InstantBlue (Expedeon) and scanned. Images were analysed using Image Lab (Bio-Rad) software. For mass spectrometry analyses, we carried out the same protocol but using non-radioactive ATP. The samples were directly loaded on an 12% SDS PAGE to be analyzed on a mass spectrometer as described below.

### Phoshorylation assays using cell extracts

About 50 x 10^6^ cells cultured in glucose-free media and supplemented with 10mM Galactose for 4 h were resuspended in kinase buffer (10mM TRIS-HCL, pH 6.7, 10mM KCl, 10mM MgCl2, 1mM DTT, 1% DDM). We then added 50 µl of purified ECSIT_CTER_ at 4 mg/ml, 10 µl of phosphatase/protease inhibitors (Protease and Phosphatase Inhibitor Cocktail, EDTA-free, Abcam) and 5 mM ATP, all processed on ice. The incubation samples were then sonicated for 2 s and and kept on ice for 5 min. We repeated this process twice. The samples were then incubated at 30 °C for 20 min to start the phosphorylation reaction and the reaction stopped by adding 50 µl of 4X SDS PAGE loading buffer (0.4% bromophenol blue, 0.4 M DTT, 0.2 M TRIS pH 6.8, 8% SDS, 40% glycerol) to each sample. The samples were directly loaded on an 12% SDS PAGE to be analyzed on a mass spectrometer as described below.

### Mass spectrometry In-gel digestion

The mass spectromety analyses were carried out at the Proteomics Core Facility from the EMBL Heildelberg. Bands corresponding to the protein of interest were cut from the gel and in-gel digestion with trypsin (Promega) was performed essentially as described in ^41^. Peptide extraction was done by sonication for 15 minutes, followed by centrifugation and supernatant collection. A solution of 50:50 water: acetonitrile, 1 % formic acid was used for a second extraction. The supernatants of both extractions were pooled and dried in a vacuum concentrator. Peptides were dissolved in 10 µL of reconstitution buffer (96:4 water: acetonitrile, 1% formic acid) and analyzed by LC-MS/MS.

### Mass spectrometry measurements

For LC-MS/MS measurement, an Orbitrap Fusion Lumos instrument (Thermo) coupled to an UltiMate 3000 RSLC nano LC system (Dionex) was used. Peptides were concentrated on a trapping cartridge (µ-Precolumn C18 PepMap 100, 5µm, 300 µm i.d. x 5 mm, 100 Å) with a constant flow of 0.05% trifluoroacetic acid in water at 30 µL/min for 6 minutes. Subsequently, peptides were eluted and separated on the analytical column (nanoEase™ M/Z HSS T3 column 75 µm x 250 mm C18, 1.8 µm, 100 Å, Waters) using a gradient composed of Solvent A (3% DMSO, 0.1% formic acid in water) and solvent B (3% DMSO, 0.1% formic acid in acetonitrile) with a constant flow of 0.3 µL/min. The percentage of solvent B was stepwise increased from 2% to 6% in 6 min, to 24% for a further 41 min, to 40% in another 5 min and to 80% in 4 min. The outlet of the analytical column was coupled directly to the mass spectrometer using the nanoFlex source equipped with a Pico-Tip Emitter 360 µm OD x 20 µm ID; 10 µm tip (MS Vil). Instrument parameters: spray voltage of 2.4 kV; positive mode; capillary temperature 275°C; Mass range 375-1650 m/z (Full scan) in profile mode in the Orbitrap with resolution of 120000; Fill time 50 ms with a limitation of 4e5 ions. Data dependent acquisition (DDA) mode, MS/MS scans were acquired in the Orbitrap with a resolution of 30000, with a fill time of up to 86 ms and a limitation of 2e5 ions (AGC target). A normalised collision energy of 34 was applied (HCD). MS2 data was acquired in profile mode.

### Mass spectrometry data processing

The raw mass spectrometry data was processed with MaxQuant (v1.6.17.0)^42^ and searched against the uniprot-proteome UP000000625 (E.coli, 4450 entries, May 2022) database containing the sequence of the protein of interest and common contaminants. The data was searched with the following modifications: Carbamidomethyl (C) as fixed modification, acetylation (Protein N-term), oxidation (M) and phosphorylation (STY) as variable modifications. The default mass error tolerance for the full scan MS spectra (20 ppm) and for MS/MS spectra (0.5 Da) was used. A maximum number of 3 missed cleavages was allowed. For protein identification, a minimum of 2 unique peptides with a peptide length of at least seven amino acids and a false discovery rate below 0.01 were required on the peptide and protein level. Match between runs was used with default parameters. The msms.txt output file of MaxQuant was used for MS1 Filtering with Skyline (v21.1.0.278)^43^.

## ACKNOWLEDGEMENTS

We are grateful to Dr. C. Mas (IBSG Grenoble) for support with ITC and mass photometry. We thank Dr. S. Bohic (ESRF, INSERM) and Dr. C. Petosa (IBS Grenoble) for discussions. We thank Dr. N. Burgess-Brown (Oxford University) for the VLCAD plasmid and Dr. P. Aloy (IRB Barcelona) for H4 cell lines. This work used the EMBL Proteomics Core Facility and the platforms of the Grenoble Instruct Center (ISBG; UMS 3518 CNRS-CEA-UJF-EMBL) with support from FRISBI (ANR-10-INSB-05-02) and GRAL (ANR-10-LABX-49-01) within the Grenoble Partnership for Structural Biology (PSB). The IBS EM facility is supported by the Rhône-Alpes Region, the Fondation Recherche Medicale (FRM), the fonds FEDER and the GIS-Infrastrutures en Biologie Sante et Agronomie (IBISA). We acknowledge the ESRF for provision of beam time on CM01. The ESRF in-house Research Program supported this work. The EM work was funded by the European Union’s Horizon 2020 research and innovation programme under grant agreement No. 647784 to IG.

## SUPPLEMENTAL INFORMATION

**Supplementary Figure 1.**
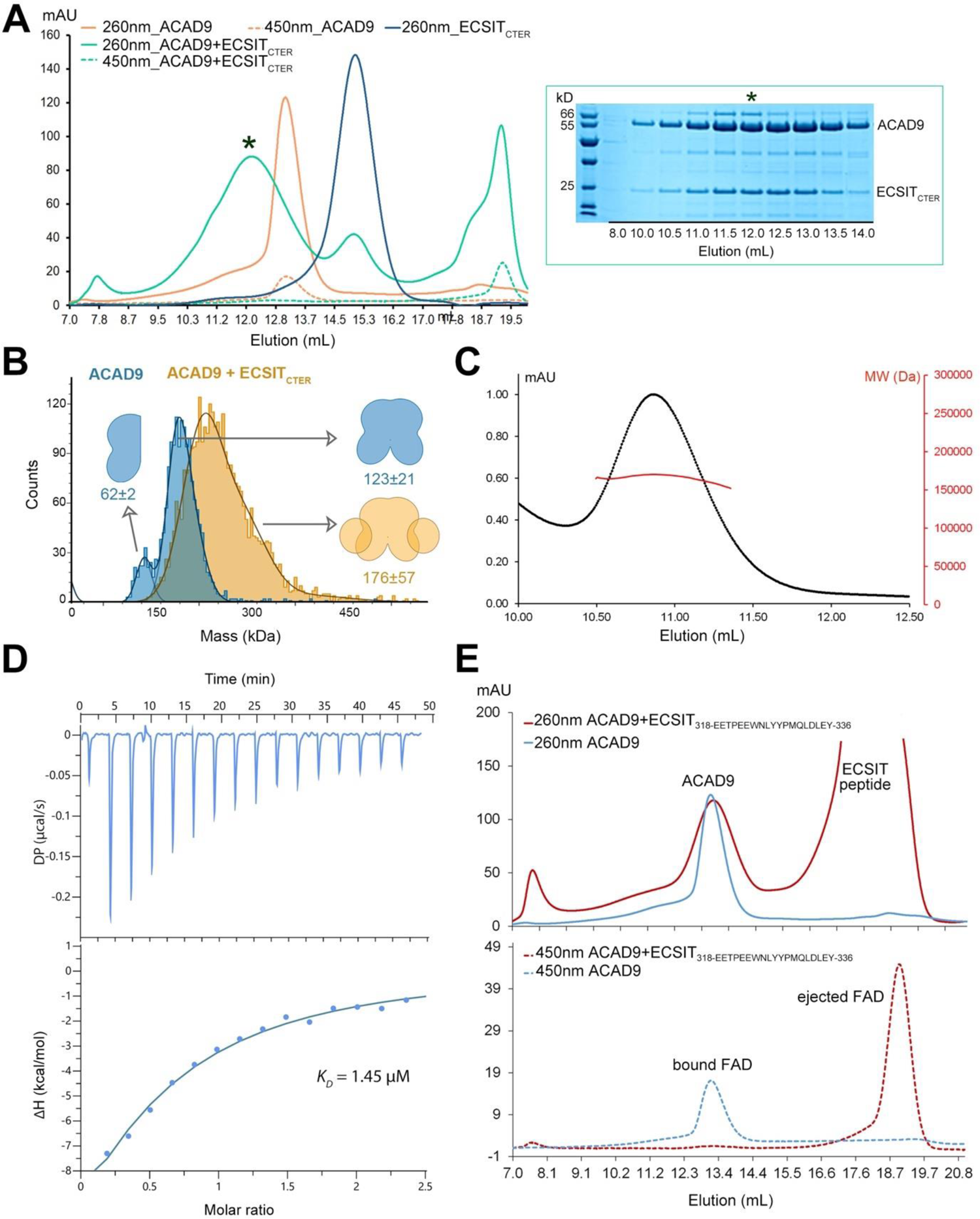
Biophysical characterisation of the ACAD9-ECSIT_CTER_ complex. **(A)** *Left*, SEC elution profiles of ACAD9 and ECSIT_CTER_ alone and in complex monitored using UV wavelengths of 260 nm and 450 nm. ACAD9_WT_ elutes at ∼13 mL (solid orange line) with FAD bound (dotted orange line); ECSIT_CTER_ elutes ∼15.3 mL. The ACAD9-ECSIT_CTER_ complex elutes ∼12mL as indicated by a * on the 260 nm signal. The corresponding 450 nm signal (dotted green line) indicates there is no FAD present in the ACAD9-ECSIT_CTER_ complex and instead is eluted much later as an unbound FAD molecule ∼19 mL. *Right*, SDS-PAGE gel showing the eluted fractions corresponding to the ACAD9-ECSIT_CTER_ complex. The * corresponds to the apex of the ACAD9-ECSIT_CTER_ elution peak as shown in the chromatogram. **(B)** Mass photometer analysis shows the primary species in the ACAD9 sample (blue) is the ACAD9 dimer with a MW of ∼123±21 kDa, with an additional smaller peak corresponding to the ACAD9 monomer species with a MW of 62±2 kDa in much lower proportion. A shift in MW is seen in the formation of the ACAD9-ECSIT_CTER_ complex, resulting in an estimated MW of 176±57, corresponding to a dimer of ACAD9 and two ECSIT_CTER_ monomers, as described in the schematic (orange). **(C)** Analysis of the ACAD9-ECSIT_CTER_ complex by SEC coupled with multi-atomic light scattering (MALS). The elution profile was monitored by excess refractive index (left axis, red). The estimated atomic mass of the complex is given by the black line, indicating that the MW of the ACAD9-ECSIT_CTER_ complex is 170 kDa, measured on the right axis. **(D)** ITC binding assay for the binding affinity between ACAD9 and ECSIT_CTER_. The equilibrium dissociation constant (K_D_) of the ACAD9-ECSIT_CTER_ complex is 1.45 μM from three replicate experiments. **(E)** SEC elution profiles of ACAD9 alone and reconstituted with the ECSIT peptide (residues 318-336), monitored using UV wavelengths of 260 nm and 450 nm. The SEC elution profile for ACAD9 (blue) shows the elution around 13 mL (top panel) with the corresponding FAD signal eluting at the same volume (blue, bottom panel), indicating FAD-bound ACAD9. When reconstituted with the ECSIT peptide, the elution profile (red, top panel) shows no significant shift from the ACAD9 alone profile (blue) however there is clearly a loss of FAD from the ACAD9 dimer (red, bottom panel), induced by the ECSIT peptide.

**Supplementary Figure 2.**
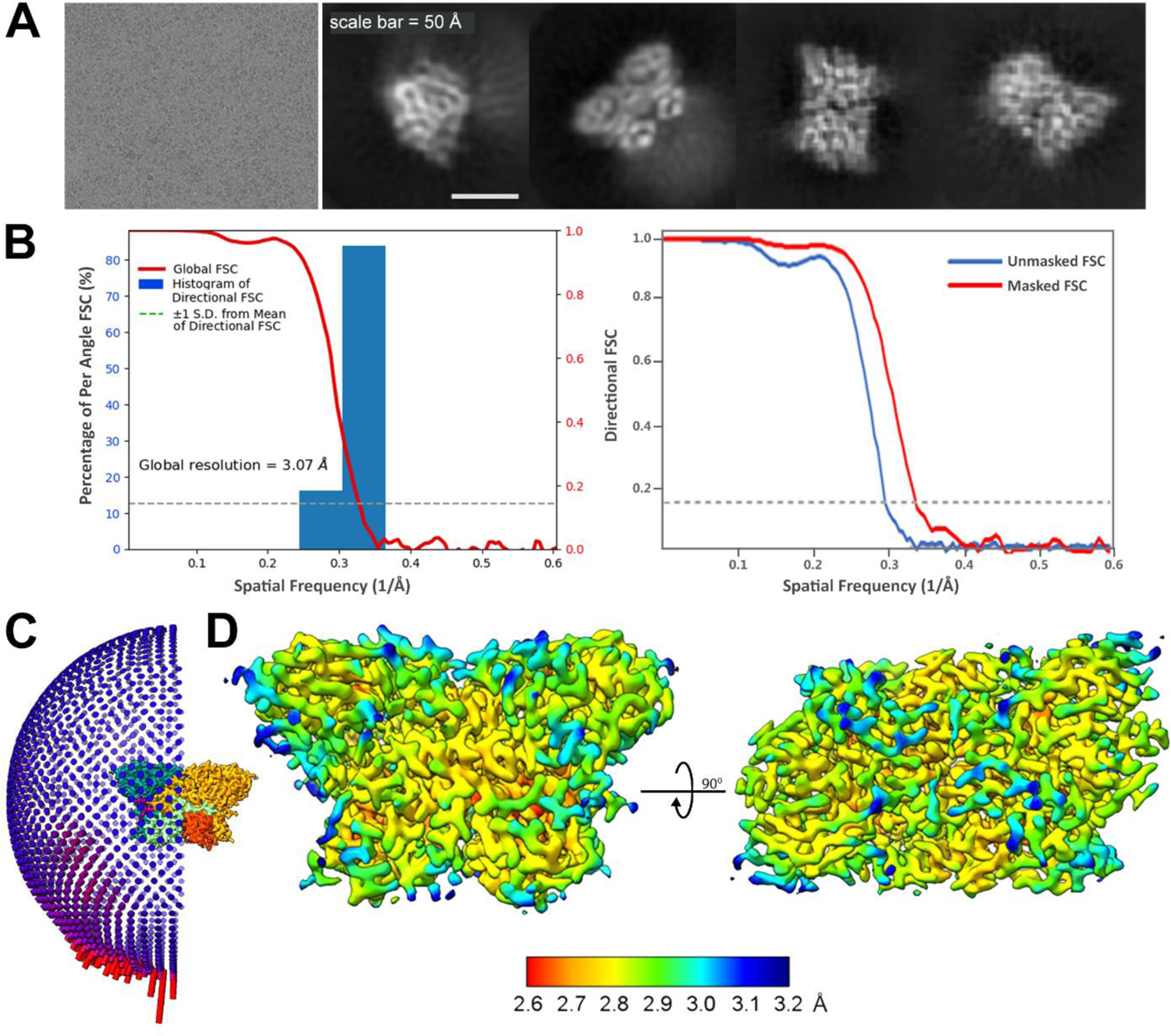
Data quality assessment of the ACAD9-ECSIT_CTER_ dataset. **(A)** Typical micrograph representative of the ACAD9-ECSIT_CTER_ dataset and selected 2D class averages. **(B)** Histogram of the directional FSC (*left*), plotting the global half-map FSC (solid red line) and where the blue bars are a histogram of 100 such values sampled evenly over the 3D FSC, indicating a global resolution of 3.07 Å. The FSC curves (right) for both the unmasked (blue) and masked (red) maps are shown, the 0.143 cut-off is represented by a grey dotted line. **(C)** Angular distribution of the particles used for the final reconstruction of ACAD9-ECSIT_CTER_. **(D)** Local resolution distribution of the final map as calculated in CryoSPARC, showing front and top particle views.

**Supplementary Figure 3:**
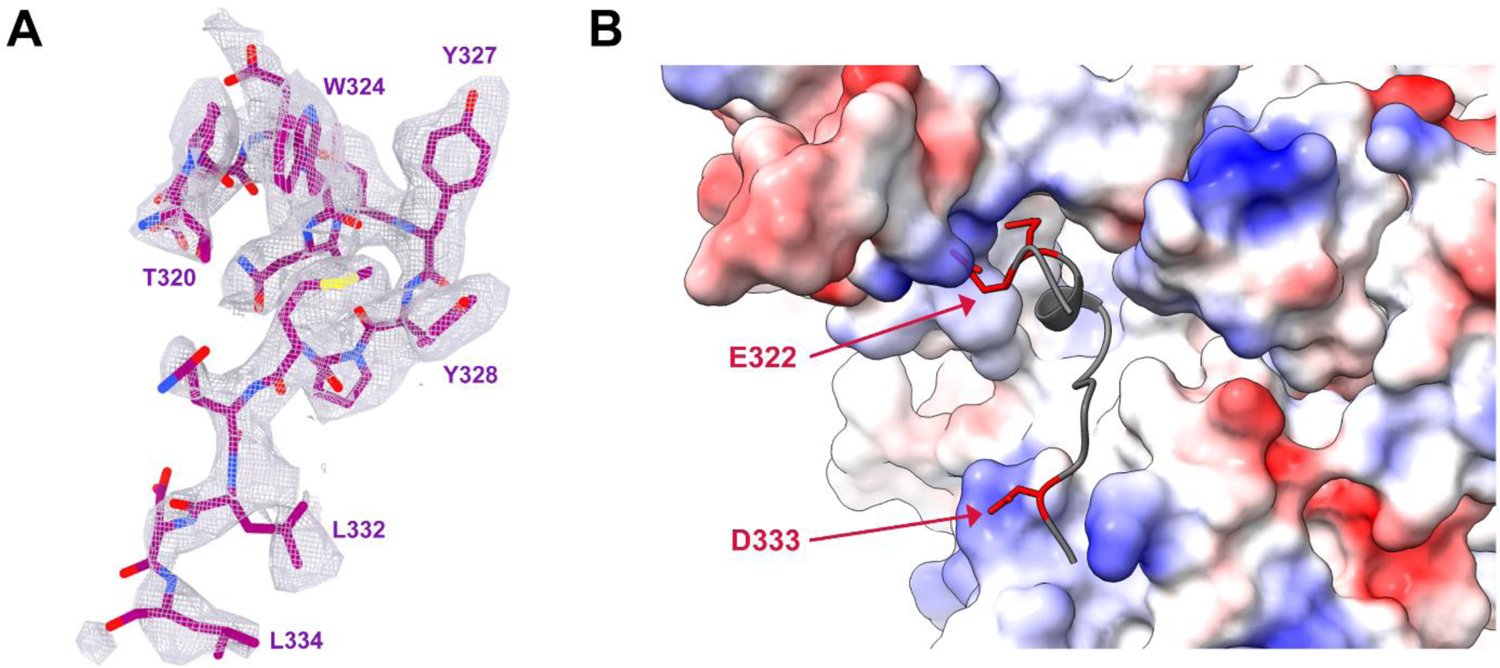
Zoom into the cryo-EM map of ACAD9-ECSIT_CTER._ **(A)** Zone of the cryo-EM map corresponding to the 15-residue peptide spanning residues 320-334 of ECSIT_CTER_ (Fig. 1). **(B)** Representation of the electrostatic properties at the ACAD9-ECSIT_CTER_ binding interface as calculated by APBS electrostatics. The surface of the junction between the vestigial and dehydrogenase domains of ACAD9 is mainly positively charged (blue). In comparison, the ECSIT sequence modelled from the cryo-EM map contains hydrophobic and polar residues (grey) but principally carries a negative charge. Negatively charged residues Glu322, Glu323 and Asp333 are represented as red sticks for clarity.

**Supplementary Figure 4:**
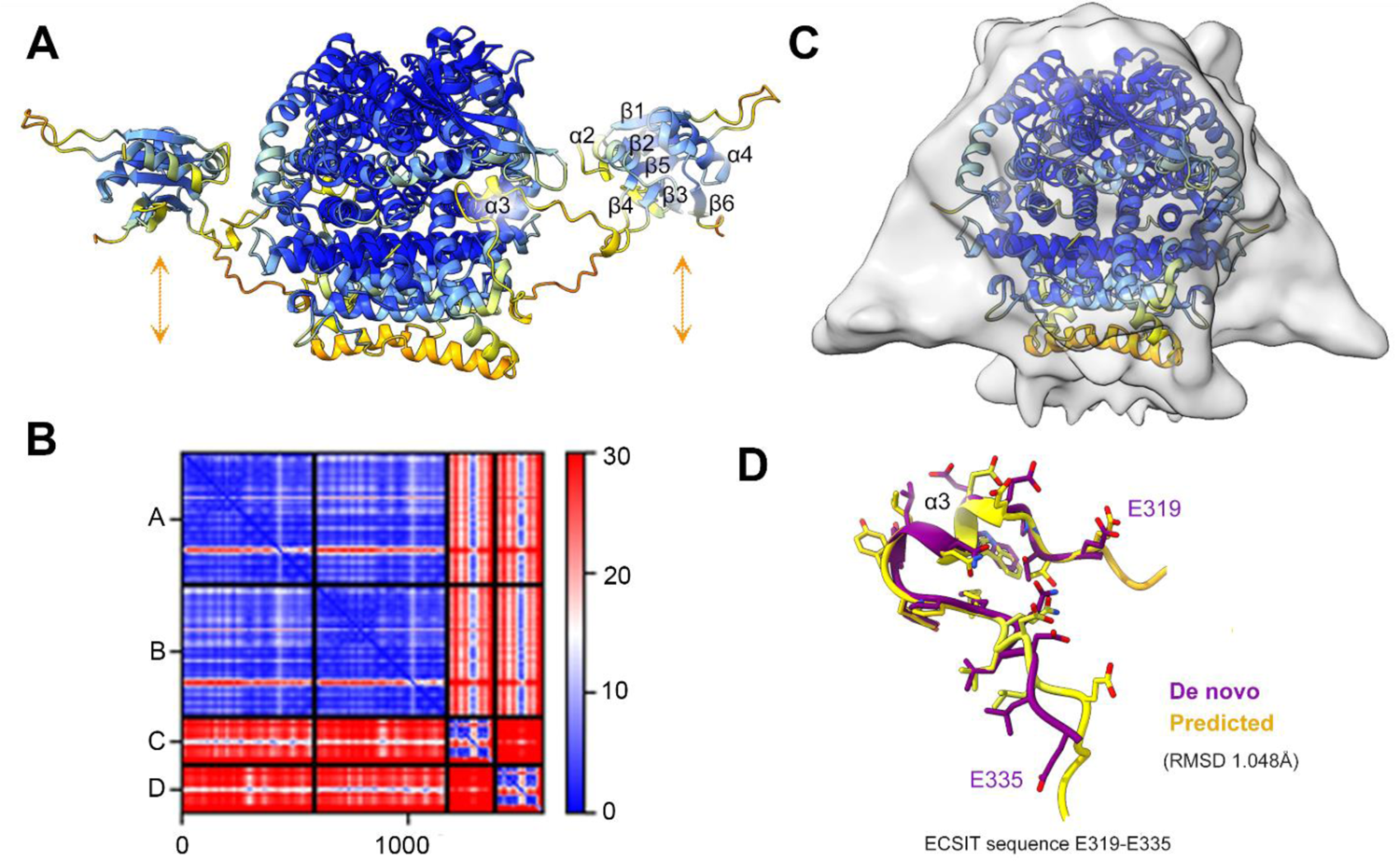
AlphaFold2 (AF) calculations predict the same ECSIT binding as that observed in our experimental model. **(A)** Calculated model from AF2 showing the dimer of ACAD9 and two ECSIT_CTER_ monomers from a side view. The docked model is coloured according to the pLDDT score, indicating a per-residue estimate of confidence. pLDDT scores are coloured as follows: pLDDT < 50 (very low) = orange, 70 > pLDDT >=50 (low) = yellow, 90 > pLDDT >=70 (confident) = light blue, pLDDT > 90 (high) = dark blue. The colour of the ECSIT binding loop (residues 320-334 corresponding to the β3-β4 loop in the AF2 model) are coloured yellow, indicating low per-residue confidence, however the Predicted Aligned Error (PAE) plot shown in **(B)** indicates a low level of error, indicating that while the prediction has a low pLDDT score of its confidence in the orientation of the β3-β4 loop that binds to ACAD9, the low PAE error suggests that the prediction is confident of their relative positions within the complex. The y-axis indicates A and B correspond to the first and second ACAD9 monomers and C and D correspond to the two ECSIT monomers. The x-axis gives the residue number from A -> D across the whole complex. The PAE scores between 0 (blue, low error) and 30 (red, high error). **(C)** Although in our earlier low-resolution map of ACAD9-ECSIT_CTD_ we could only docked the ACAD9 homodimer^11^, at low threshold it shows a density extension that appears to be very closed to the ECSIT β3-β4 loop flexibly connected to strands β3 and β4 of the RNAseH-like domain, suggesting that this domain could be positioned within the extra density with only a minor adjustment of the long predicted β3-β4 loop, as proposed by the orange arrows in **(A)**. **(D)** An alignment of the ECSIT loop from the AF2 prediction (yellow) with our experimental model (purple) shows high agreement (RMSD = 1.048 Å).

**Supplementary Figure 5.**
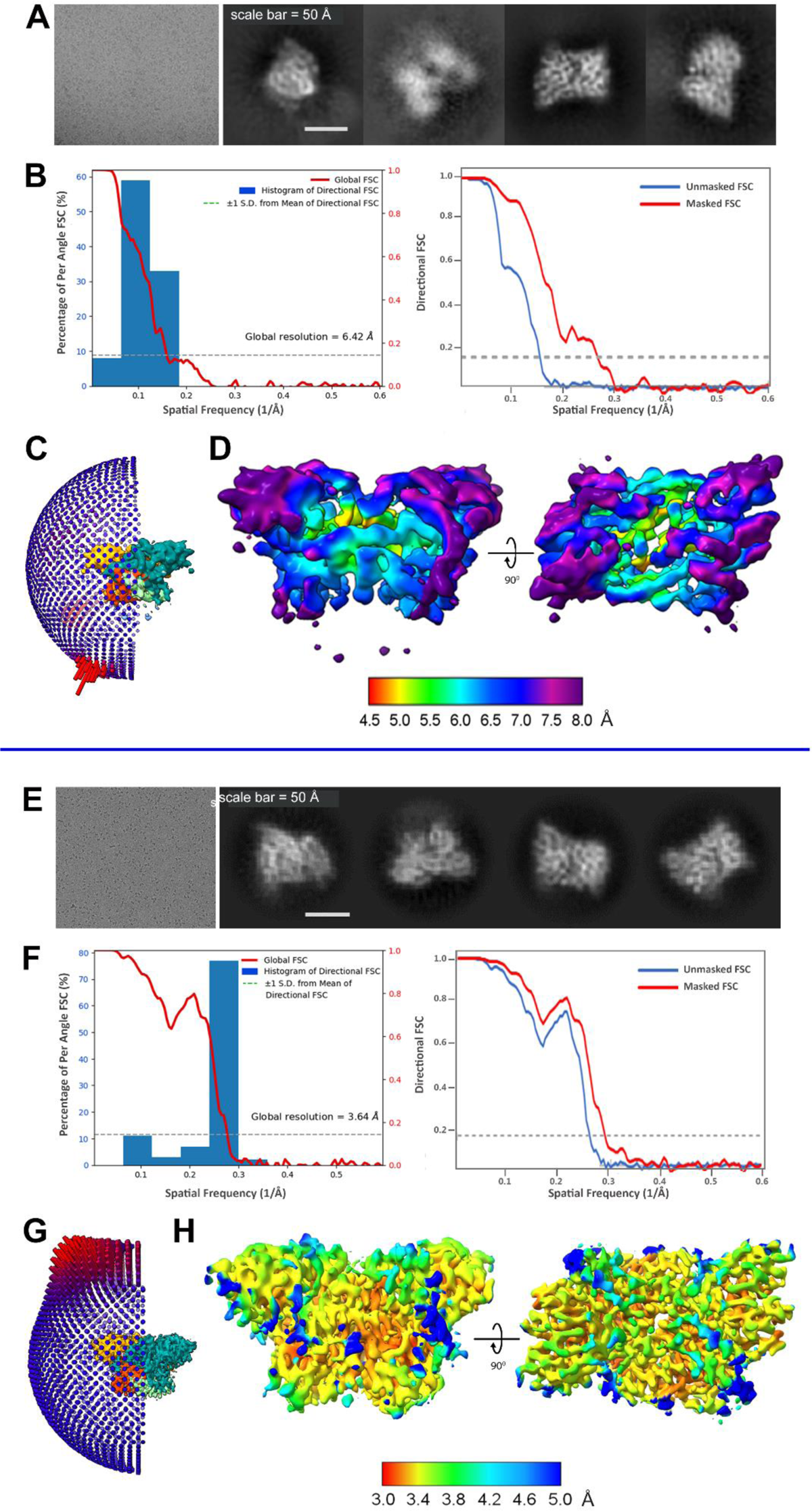
Data quality assessment of the ACAD9_WT_ and ACAD9_S191A_ datasets. **(A)** Typical micrograph representative of the ACAD9_WT_ dataset and selected 2D class averages of the ACAD9_WT_ particle. **(B)** Histogram of the directional FSC (*left*), indicating a global resolution of 6.42 Å. The FSC curves (right) for both the unmasked (blue) and masked (red) maps are shown, the 0.143 cut-off is represented by a grey dotted line. **(C)** Angular distribution of the particles used for the final reconstruction of ACAD9_WT_. **(D)** Local resolution distribution of the final ACAD9_WT_ map as calculated in CryoSPARC, showing front and top particle views. The local resolution is higher in the core of the protein (∼4.0 Å) whereas the solvent accessible areas of the protein are less well resolved (∼8.0 Å). **(E)** Typical micrograph representative of the ACAD9_S191A_ dataset and selected 2D class averages of the ACAD9_S191A_ particle. **(F)** Histogram of the directional FSC (*left*), indicating a global resolution of 3.64 Å. The FSC curves shown as in B. **(G)** Angular distribution of the particles used for the final reconstruction of ACAD9_S191A_. **(H)** Local resolution distribution of the final ACAD9_S191A_ map, showing front and top particle views.

**Supplementary Figure 6.**
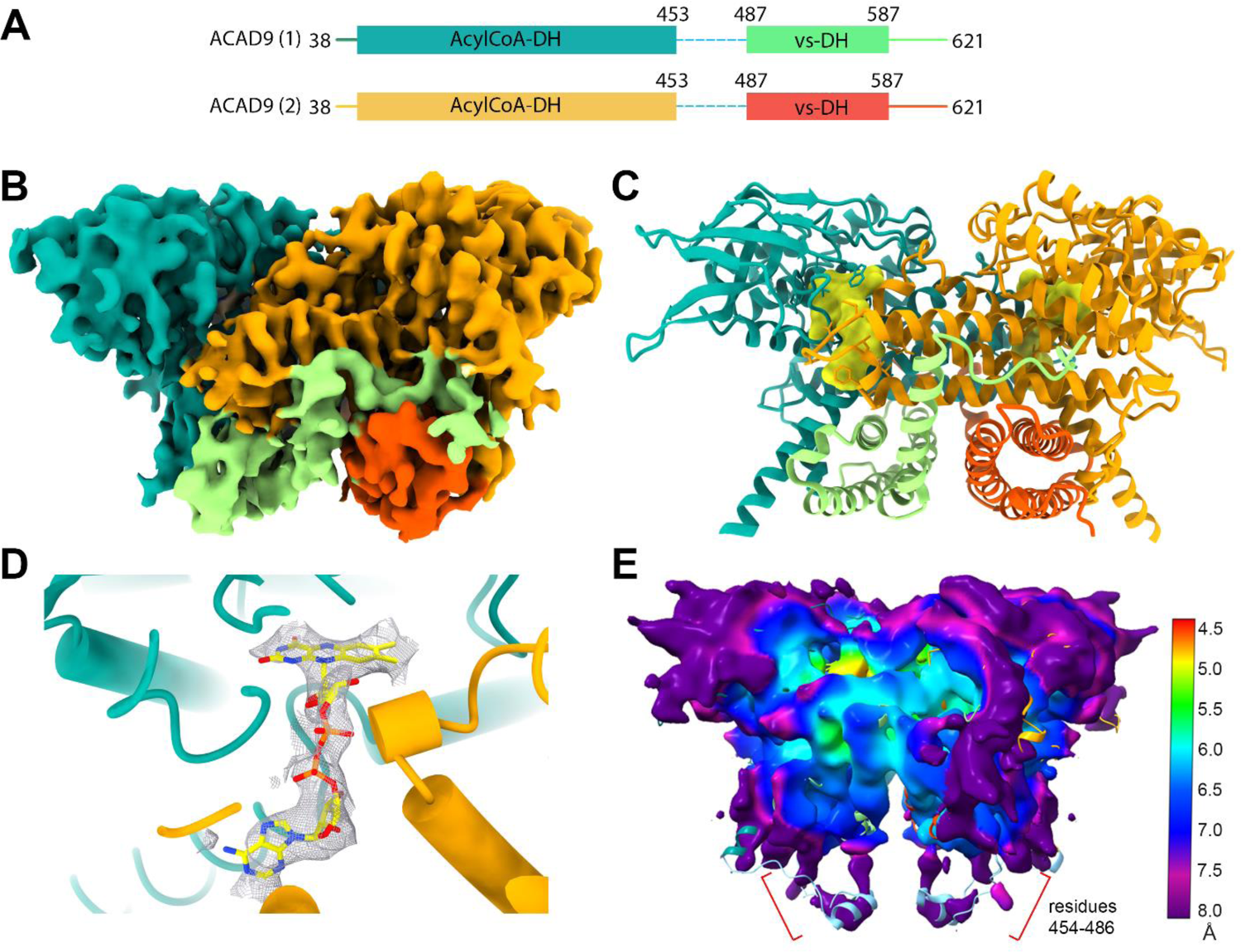
Cryo-EM structure of ACAD9 alone. **(A)** Cryo-EM map of ACAD9_S191A_ viewed from the front, coloured as in Fig. 1. **(B)** Final model of ACAD9_S191A_ with the density of the FAD highlighted in yellow. **(C)** Close-up of the FAD density in the ACAD9_S191A_ cofactor pocket, confirming that ACAD9_S191A_ is in the active Acyl-CoA dehydrogenase form of ACAD9. **(D)** The final reconstruction of ACAD9_WT_ depicted as a local resolution map. Reducing the threshold enables visualization of faint densities at the base of the protein, close to the location of the unmodelled residues 454-486. The homology model was modified to fit these residues (light blue) into the low-resolution density, showing that these previously unmodelled residues are located close to the base of the ACAD9 dimer.

**Supplementary Figure 7.**
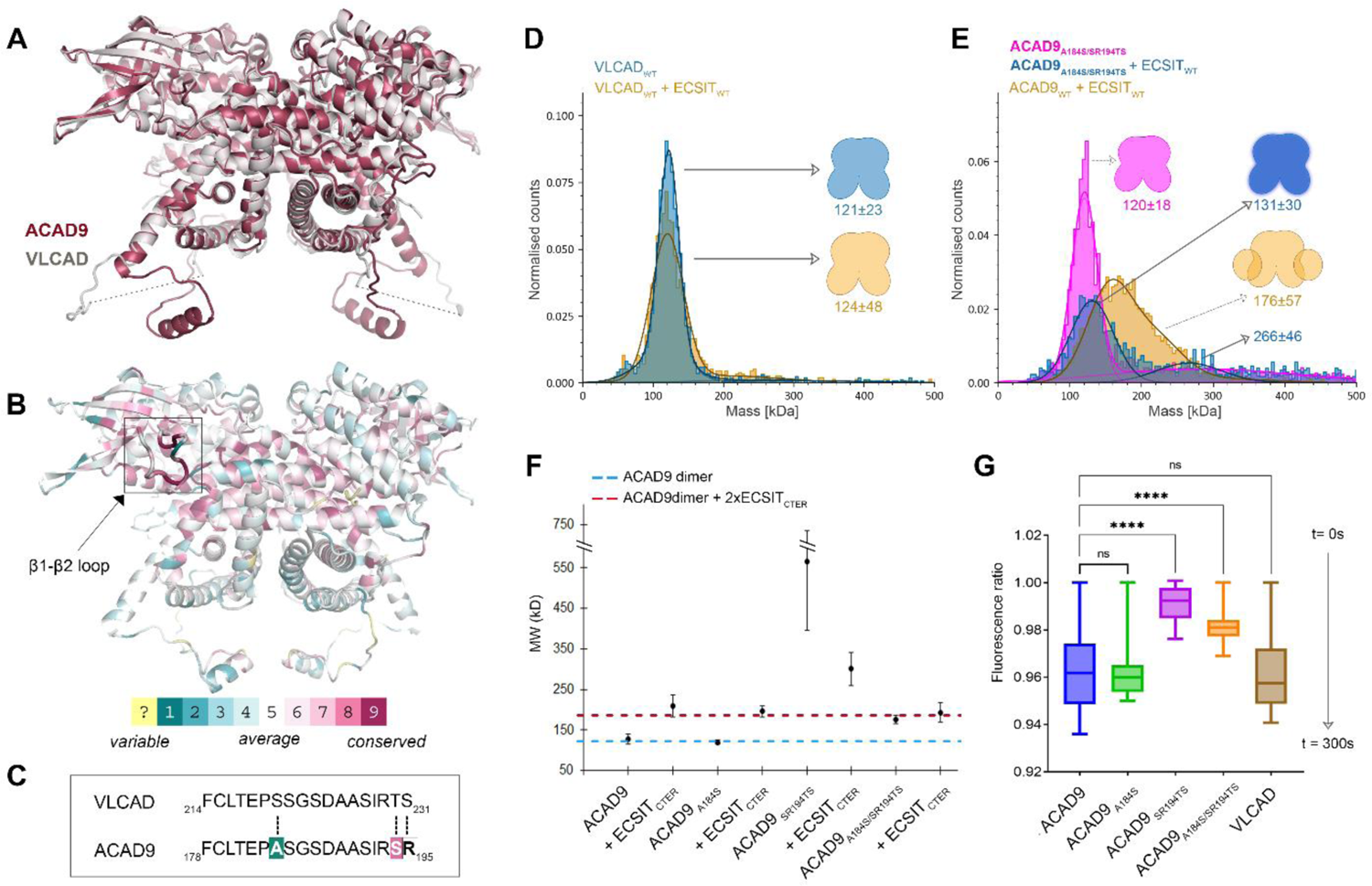
Conformational analysis of the ACAD9 gatekeeper loop based on VLCAD-derived mutants. **(A)** Superposition of VLCAD crystal structure and the ACAD9 cryo-EM structure**. (B)** Representation of ACAD9 structure based on the evolutionary conserved residues estimated by the ConSurf prediction software**. (C)** The sequences of ACAD9 and VLCAD when aligned with the loop present on ACAD9 that has been found to undergo a large conformational change upon ECSIT binding. The residues that were mutated in ACAD9 to mimic the behaviour of VLCAD are shown in bold. **(D)** Mass photometer data of VLCAD (*left*) shows the formation of a very stable dimeric species (blue) with a MW of 121±23 kDa and when reconstituted with ECSIT_CTER_ no shift towards higher MW is seen (orange), MW estimation 124±48 kDa, confirming that VLCAD does not form a complex with ECSIT_CTER_. In contrast, when reconstituted together ACAD9 and ECSIT_CTER_ form a stable complex with a MW of 176±57 kDa (*right*, orange). **(E)** When the three residues on the ACAD9 loop are mutated to mimic VLCAD, ACAD9 forms a dimer (pink) with a MW = 120±18 kDa but with some higher order species (MW = 282±127 kDa), and upon reconstitution with ECSIT (blue), a proportion of the protein remains dimeric (MW = 131±30 kDa) with some higher order species (MW = 266±46kDa). Indicating that the double mutant decreases the stability of the complex in more VLCAD-like behaviour, however, is able to retain the ability to bind to ECSIT. **(F)** DLS measurements of ACAD9 mutants designed to mimic the loop of VLCAD. The ACAD9_A184S_ mutant exhibits similar behaviour to that of ACAD9_WT_ both alone and in complex with ECSIT_CTER_. The double mutant ACAD9_SR194TS_ has a large number of higher order species present, indicating that this mutation destabilises ACAD9, although the effect is not as profound in complex with ECSIT_CTER_; some complex is formed but at higher MW than the ACAD9_WT_-ECSIT_CTER_. In contrast, the double mutant ACAD9_A184S/SR194TS_ shows no change in MW between ACAD9_A184S/SR194TS_ alone and after reconstitution of ECSIT, however when combined with the mass photometer data we can attribute this to a reduction in protein stability however the ability to bind ECSIT_CTER_ is retained. **(G)** Acyl-CoA dehydrogenase (ACAD) activity of ACAD9_WT_, ACAD9 single and double mutants and VLCAD as determined by an ETF fluorescence reduction assay. After addition of the ACAD specific substrate palmitoyl-CoA (C16:0), there is a clear loss of ETF fluorescence in ACAD9_WT_, ACAD9_A184S_ and VLCAD in comparison to the other ACAD9 mutants, over 300 s of reaction measurement.

**Supplementary Figure 8.**
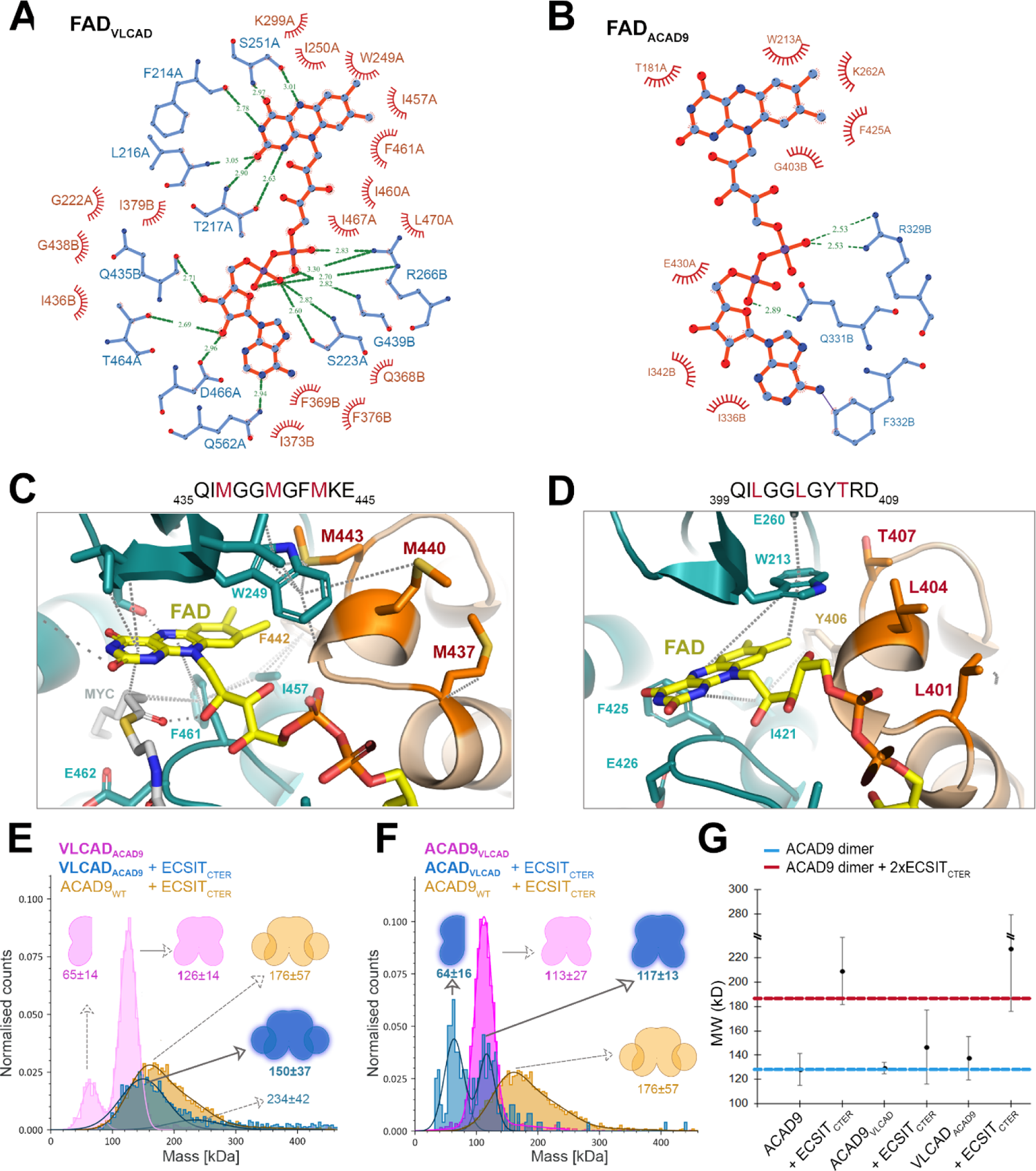
The architectures of the FAD binding pockets are different between ACAD9 and VLCAD. The FAD binding pocket of VLCAD is organised to bind and retain FAD more tightly than the FAD binding pocket of ACAD9. **(A)** Ligplot Analysis of the bonding interactions between the FAD cofactor and the VLCAD protein. The high number of bonds formed between the protein and cofactor suggests that the FAD molecule is securely held in place. For FAD ejection to occur in this case, a large number of interactions would need to be perturbed to allow the FAD molecule to freely exit the protein. **(B)** Ligplot analysis of the bonding interactions between the FAD cofactor and the ACAD9 protein. In comparison to VLCAD, a reduced number of bonding interactions are present in the FAD-bound ACAD9 protein. This may be related to why ACAD9 has the propensity to lose FAD upon ECSIT binding whereas VLCAD cannot. **(C)** In VLCAD, a conserved Arginine (R326) forms multiple hydrogen bonds to the pyrophosphate moiety of the FAD molecule. Three Methionine residues (M403, M400, M397) are situated at the top of the VLCAD FAD site and contribute to the stability of the FAD molecule by M400 and M403 forming sulphur-π interactions with the isoalloxazine moiety and W209. **(D)** In ACAD9, R329 is shown to interact with the pyrophosphate moiety, however, it does not form two bonds, instead a secondary bonding interaction comes from T438. In place of three Methionine residues present in VLCAD, ACAD9 has a Threonine and two Leucine residues, unable to interact with the isoalloxazine ring and W213, thus not providing stability to the FAD molecule. **(E)** Mass photometer data showing that when VLCAD is mutated to mimic ACAD9 (^437^MGGMGFM^4^^43^ to ^437^LGGLGYT^4^^43^, VLCAD_ACAD9_, pink) exists primarily as a dimer with a MW of 126±14 kDa with a small percentage existing in the monomeric form, MW 65±14 kDa, a reduction in stability in comparison to VLCAD_WT_ (Fig. S9B). When reconstituted with ECSIT_CTER_, there is a shift in the MW of VLCAD_ACAD9_, indicating the formation of a VLCAD_ACAD9_-ECSIT_CTER_ complex (shown in blue), similar to that observed for the ACAD9_WT_-ECSIT_CTER_ complex (orange). **(F)** The reverse mutation of ACAD9 to mimic VLCAD (^401^LGGLGYT^407^ to ^401^MGGMGFM^407^, ACAD9_VLCAD_) seems to produce a more stable ACAD9 (pink) exhibiting behaviour more typically associated with VLCAD_WT_. Upon reconstitution of ECSIT_CTER_, we see an appearance of a monomeric ACAD9_VLCAD_ (blue) indicating the destabilisation of the ACAD9 dimer, however no significant shift is seen to a higher MW, demonstrating a stark reduction in the ability of ACAD9_VLCAD_ to form a complex with ECSIT, in comparison to the ACAD9_WT_-ECSIT_CTER_ (orange). **(G)** DLS measurements comparing the effect of the mutants designed to reverse the behaviour of ACAD9 and VLCAD. Measurements of ACAD9_WT_, ACAD9_VLCAD_ and VLCAD_ACAD9_ show similar behaviour, formation of dimers. However, in the case of ACAD9_VLCAD_ designed to mimic the behaviour of VLCAD, there is a stark reduction in complex formation in comparison to ACAD_WT_-ECSIT_CTER_, implying that this mutation does indeed evoke VLCAD-like tendencies in its interaction with ECSIT_CTER_. The reverse is also true in the case of VLCAD_ACAD9_, a large average MW is seen, indicating the formation of a VLCAD_ACAD9_-ECSIT_CTER_ complex however with the addition of higher order species present too. This indicates that VLCAD_ACAD9_ behaves less like VLCAD_WT_ and more like ACAD9_WT_ through the mutation of these key residues.

**Supplementary Figure 9.**
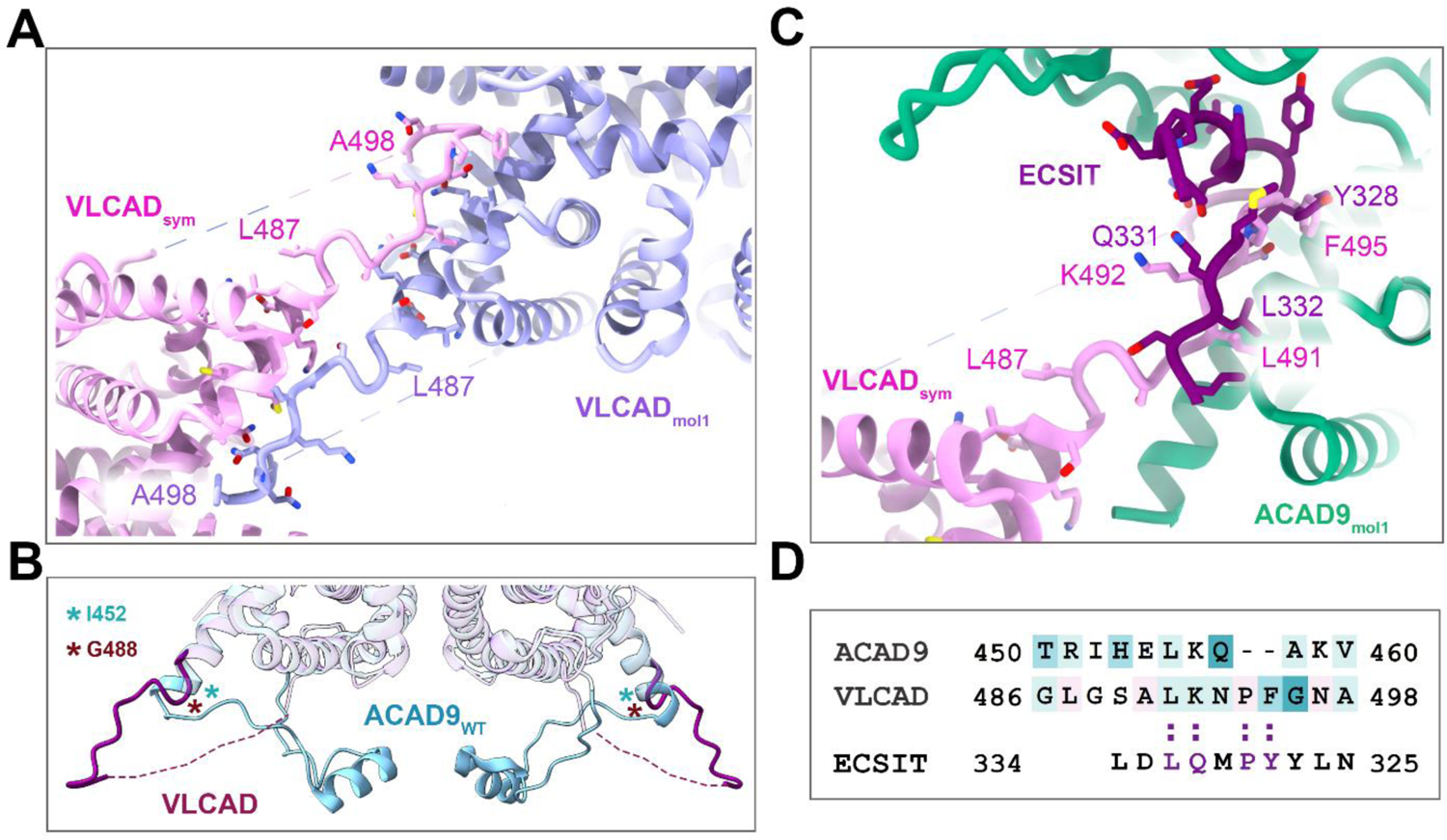
Quaternary structure of VLCAD reveals insights into the bottom loop residues and the accessibility to the ECSIT binding site in ACAD9. (A) The crystal packing of VLCAD is formed mainly by symmetric contacts involving the 485-498 loops of symmetry-related molecules. Closeup of the interaction between symmetry-related loops. (B) The structural comparison of the ACAD9_WT_ 454-486 residues with the VLCAD 490-522 residues reveal that that flexible region adopts an extended conformation in VLCAD as compared to the closed conformation observed in ACAD9_WT_ (Fig. S8C). (C) Structural alignment of the VLCAD quaternary structure and the ACAD9-ECSIT_CTER_ structure, highlighting those residues occupying similar positions in each respective model. (D) Pairwise sequence alignment of ACAD9 and VLCAD around the 35-residue region that has the greatest difference between the ACAD9 and VLCAD and that has been postulated as the determining feature for CI assembly^6^. Conserved residues between ACAD9 and VLCAD are highlighted in purple. The residues are coloured based on the ConSurf prediction (Fig. S9B), highlighting the poor conservation within ACAD9 and VLCAD in this region. Structural alignment of the VLCAD and ACDA9-ECSIT_CTER_ structures shows that the VLCAD 484-498 loop aligns well with the ECSIT sequence visible in the cryo-EM structure, suggesting that the VLCAD loop actually occupies, and perhaps blocks, the ECSIT binding site, in contrast to ACAD9.

**Supplementary Figure 10.**
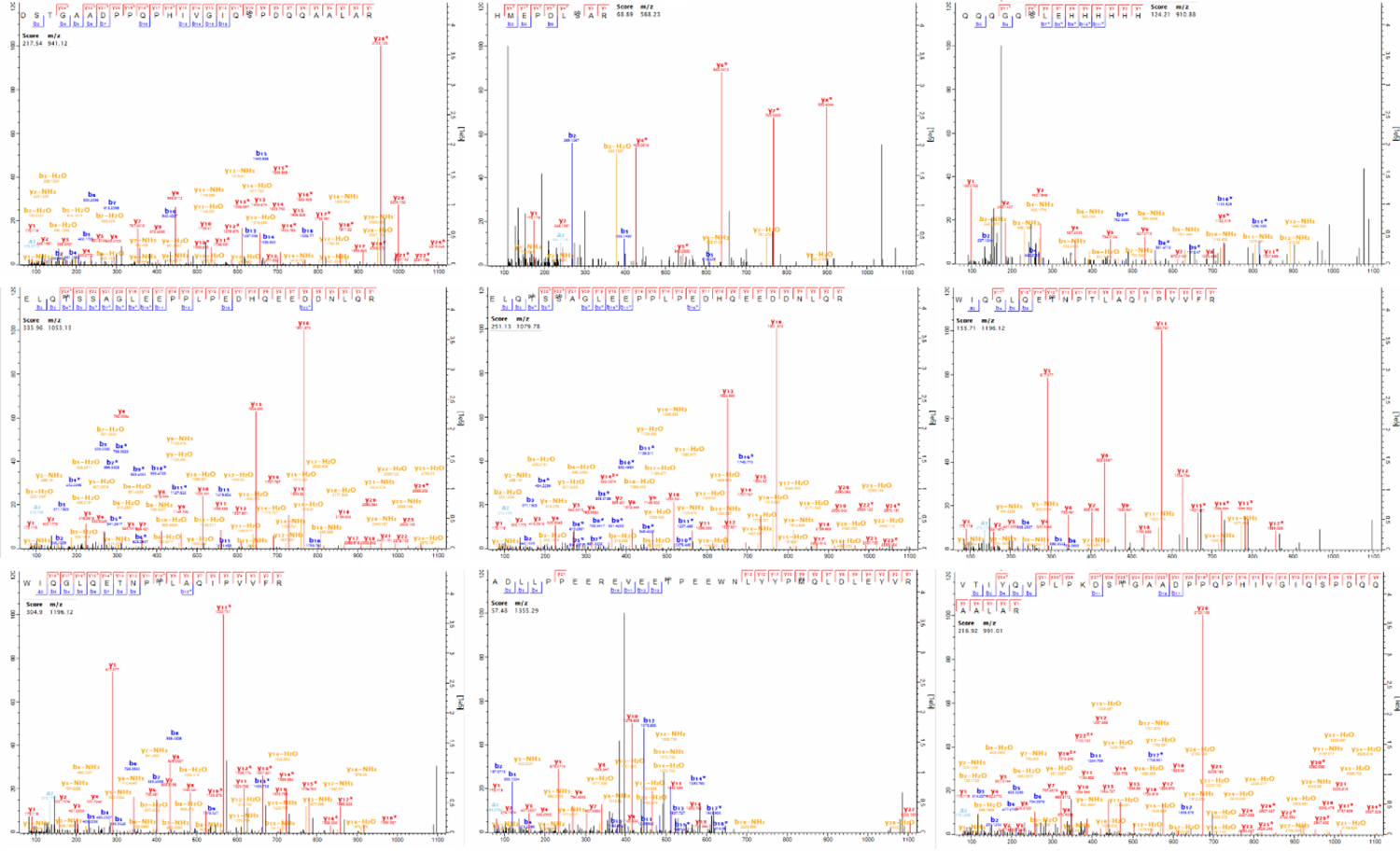
MS/MS spectra of the annotated ECSIT_CTER_ phosphosites *in vitro*. Complete sequence coverage is achieved while neutral losses are minimal, enabling unambiguous localization of the phosphosites.

**Supplementary Figure 11.**
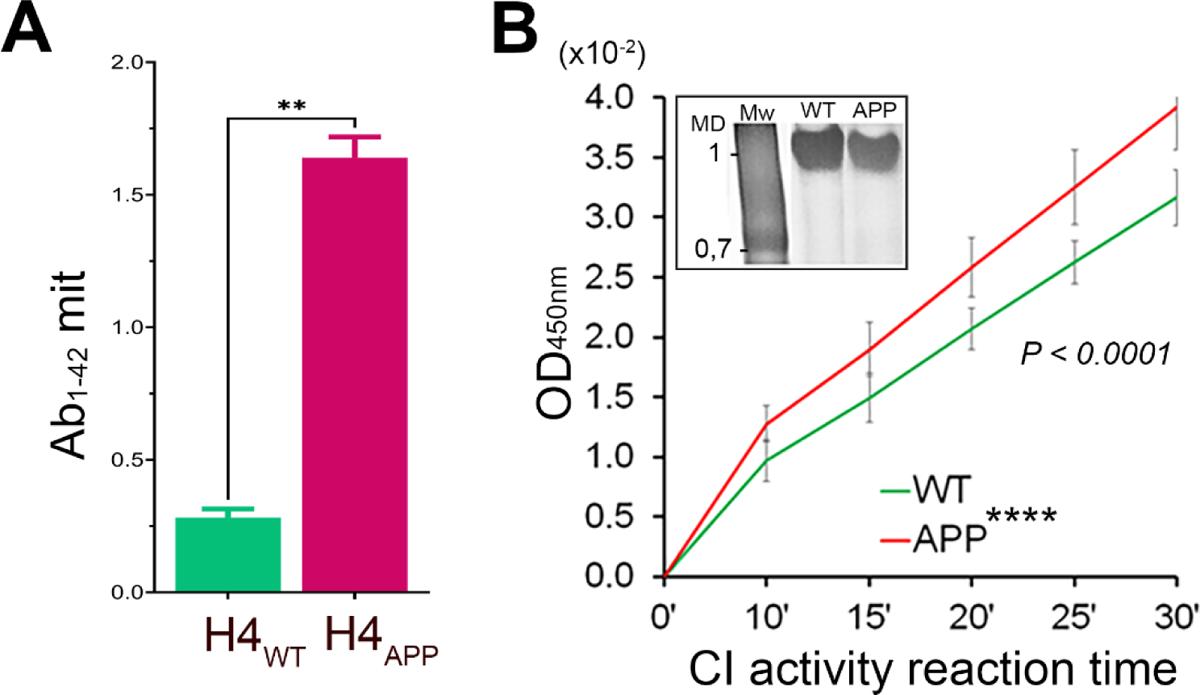
CI activity in human neuronal mitochondria and under Aβ oligomeric conditions. **(A)** Aβ_1-42_ detection in mitochondria isolated from WT and APP cells by ELISA immunoassays (n=4± SD, *\*\*P < 0.001*). H4 neuroglioma cells overexpress the human amyloid precursor protein (APP) carrying the Alzheimer’s-related Swedish mutation (KM670/671NL). **(B)** NADH-dehydrogenase activity assay of immunopurified CI in WT and APP cells (n=3± SD, \*\**\*\*P < 0.0001*). In the *inset*, native gel showing purified fully assembled CI (1 MDa) from both cell types and subjected to the activity assays.

---

**Supplementary Table 1.**
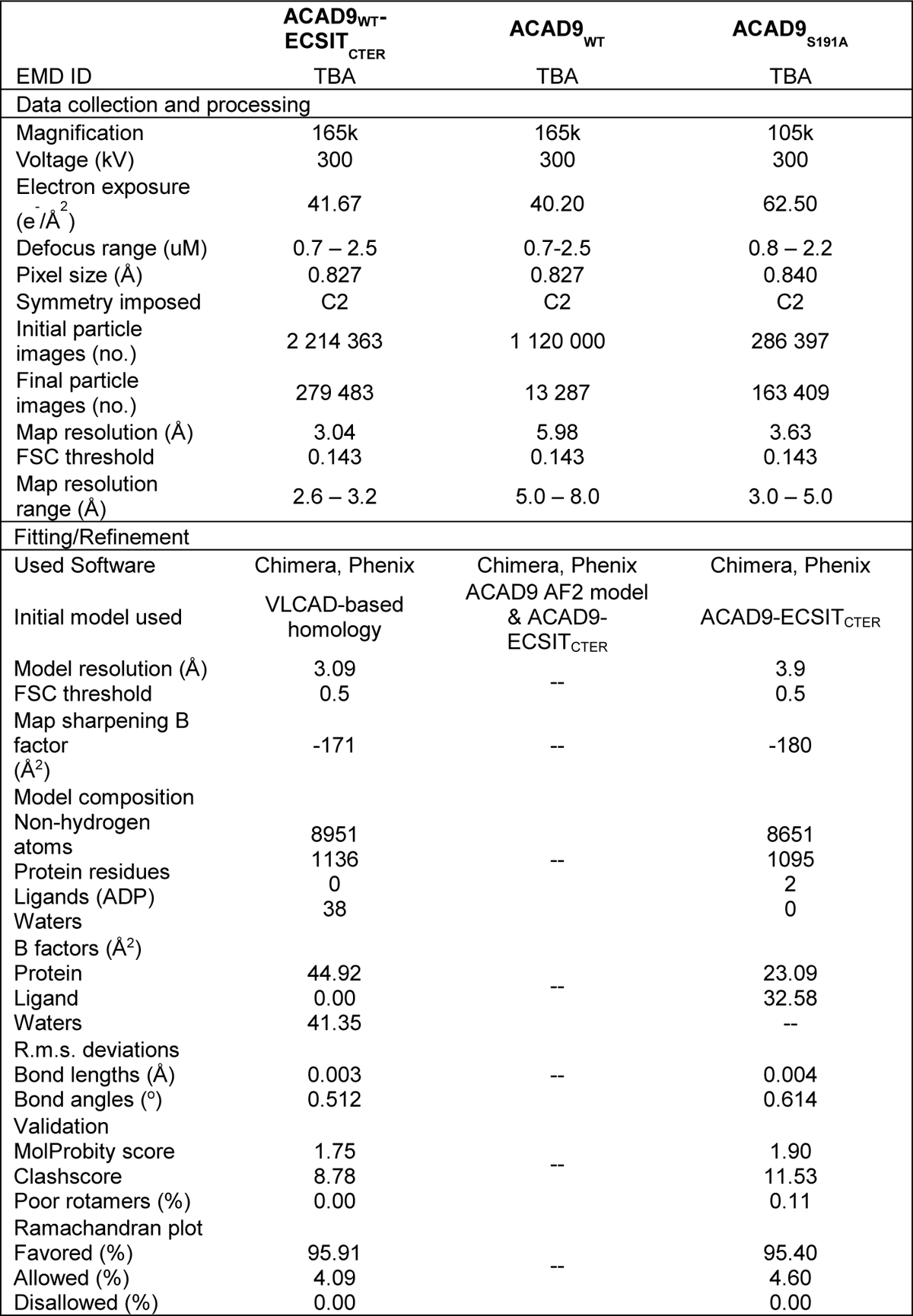
Cryo-EM data collection, processing and model refinement statistics for ACAD9-ECSIT_CTER,_ ACAD9_WT_ and ACAD9_S191A_.

**Supplementary Table 2.**
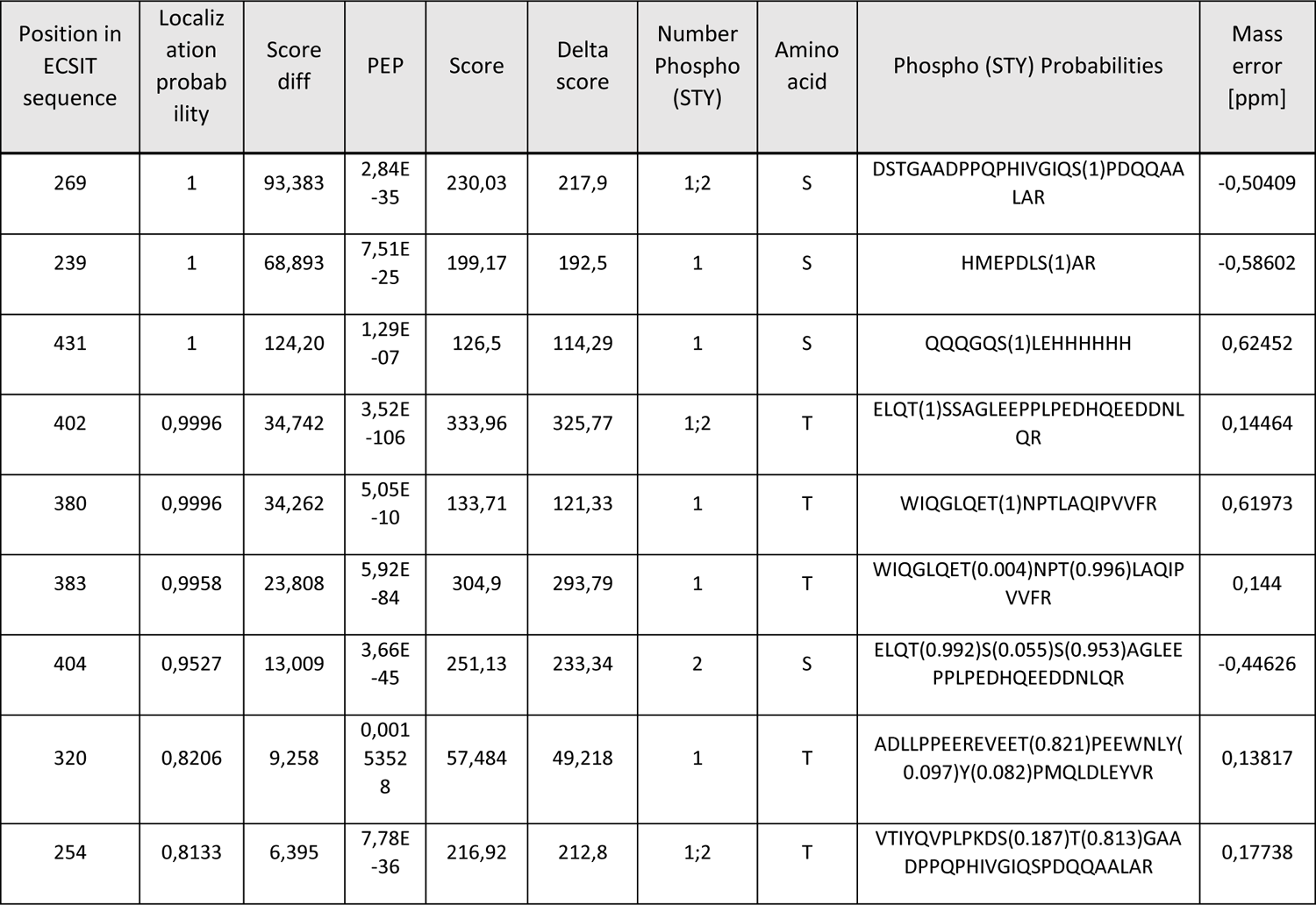
Phosphosites detected by mass spectrometry data from *in vitro* ECSIT_CTER_ phosphorylation assays. Probability of the correct localization of the modification, above or equal to 0.75 is considered as localized correctly. PEP: posterior error probability. Score: the Andromeda score of the best modified peptide. Delta score: the Andromeda delta score. Number of Phospho (STY): different numbers of phosphorylations on the given peptide. Amino acid: the amino acid which is modified. Phospho (STY) Probabilities: Amino acid sequence including localization probabilities for all considered potential sites. Charge: charge state of the precursor ion. Mass error [ppm]: mass error [ppm] of the precursor ion.

**Supplementary Table 3.**
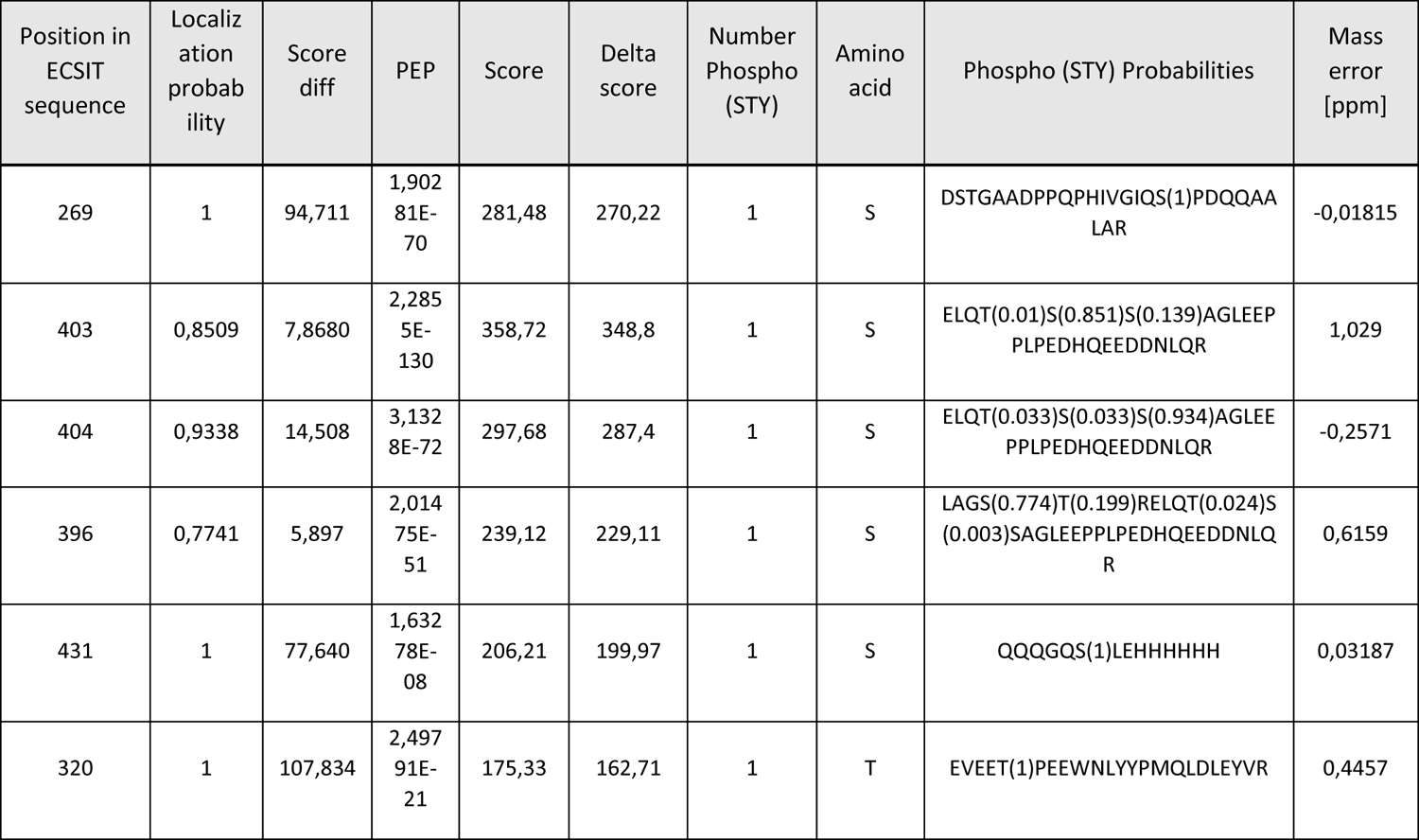
Phosphosites detected by mass spectrometry data from *ex cellulo* ECSIT_CTER_ phosphorylation assays. Probability of the correct localization of the modification, above or equal to 0.75 is considered as localized correctly. PEP: posterior error probability. Score: the Andromeda score of the best modified peptide. Delta score: the Andromeda delta score. Number of Phospho (STY): different numbers of phosphorylations on the given peptide. Amino acid: the amino acid which is modified. Phospho (STY) Probabilities: Amino acid sequence including localization probabilities for all considered potential sites. Charge: charge state of the precursor ion. Mass error [ppm]: mass error [ppm] of the precursor ion.

| Position in ECSIT sequence | Localization probability | Score diff | PEP | Score | Delta score | Number Phospho (STY) | Amino acid | Phospho (STY) Probabilities | Mass error [ppm] |
| --- | --- | --- | --- | --- | --- | --- | --- | --- | --- |
| 269 | 1 | 94,711 | 1,90281E-70 | 281,48 | 270,22 | 1 | S | DSTGAADPPQPHIVGIQS(1)PDQQAA<br>LAR | -0,01815 |
| 403 | 0,8509 | 7,8680 | 2,2855E-130 | 358,72 | 348,8 | 1 | S | ELQT(0.01)S(0.851)S(0.139)AGLEEP<br>PLPEDHQEEDDNLQR | 1,029 |
| 404 | 0,9338 | 14,508 | 3,1328E-72 | 297,68 | 287,4 | 1 | S | ELQT(0.033)S(0.033)S(0.934)AGLEE<br>PPLPEDHQEEDDNLQR | -0,2571 |
| 396 | 0,7741 | 5,897 | 2,01475E-51 | 239,12 | 229,11 | 1 | S | LAGS(0.774)T(0.199)RELQT(0.024)S<br>(0.003)SAGLEEPPLPEDHQEEDDNLQ<br>R | 0,6159 |
| 431 | 1 | 77,640 | 1,63278E-08 | 206,21 | 199,97 | 1 | S | QQQGQS(1)LEHHHHHH | 0,03187 |
| 320 | 1 | 107,834 | 2,49791E-21 | 175,33 | 162,71 | 1 | T | EVEET(1)PEEWNLYYPMQLDLEYVR | 0,4457 |

## REFERENCES

1. Moreira, P. I., Carvalho, C., Zhu, X., Smith, M. A. & Perry, G. Mitochondrial dysfunction is a trigger of Alzheimer’s disease pathophysiology. Biochimica et Biophysica Acta 1802, 2–10 (2010).

2. Zhang, L. et al. Modulation of mitochondrial complex I activity averts cognitive decline in multiple animal models of familial Alzheimer’s Disease. EBioMedicine 2, 294–305 (2015).

3. Wang, W., Zhao, F., Ma, X., Perry, G. & Zhu, X. Mitochondria dysfunction in the pathogenesis of Alzheimer’s disease: recent advances. Mol Neurodegener 15, 30 (2020).

4. Vinothkumar, K. R., Zhu, J. & Hirst, J. Architecture of mammalian respiratory complex I. Nature 515, 80–84 (2014).

5. Formosa, L. E. et al. Dissecting the Roles of Mitochondrial Complex I Intermediate Assembly Complex Factors in the Biogenesis of Complex I. Cell Reports 31, 107541 (2020).

6. Nouws, J. et al. Acyl-CoA dehydrogenase 9 is required for the biogenesis of oxidative phosphorylation complex I. Cell Metabolism 12, 283–294 (2010).

7. Zhang, J. et al. Cloning and functional characterization of ACAD-9, a novel member of human acyl-CoA dehydrogenase family. Biochem Biophys Res Commun 297, 1033–1042 (2002).

8. Kopp, E. et al. ECSIT is an evolutionarily conserved intermediate in the Toll/IL-1 signal transduction pathway. Genes Dev 13, 2059–2071 (1999).

9. Soler-Lopez, M., Zanzoni, A., Lluis, R., Stelzl, U. & Aloy, P. Interactome mapping suggests new mechanistic details underlying Alzheimer’s disease. Genome Research 21, 364–376 (2011).

10. Soler-Lopez, M., Badiola, N., Zanzoni, A. & Aloy, P. Towards Alzheimer’s root cause: ECSIT as an integrating hub between oxidative stress, inflammation and mitochondrial dysfunction. Hypothetical role of the adapter protein ECSIT in familial and sporadic Alzheimer’s disease pathogenesis. Bioessays 34, 532–541 (2012).

11. Giachin, G. et al. Assembly of The Mitochondrial Complex I Assembly Complex Suggests a Regulatory Role for Deflavination. Angewandte Chemie International Edition English 60, 4689–4697 (2021).

12. Xia, C. et al. Molecular mechanism of interactions between ACAD9 and binding partners in mitochondrial respiratory complex I assembly. iScience 24, 103153 (2021).

13. Giachin, G., Bouverot, R., Acajjaoui, S., Pantalone, S. & Soler-Lopez, M. Dynamics of Human Mitochondrial Complex I Assembly: Implications for Neurodegenerative Diseases. Frontiers Molecular Biosciences 3, 43 (2016).

14. Ensenauer, R. et al. Human acyl-CoA dehydrogenase-9 plays a novel role in the mitochondrial beta-oxidation of unsaturated fatty acids. J Biol Chem 280, 32309–32316 (2005).

15. McAndrew, R. P. et al. Structural basis for substrate fatty acyl chain specificity: crystal structure of human very-long-chain acyl-CoA dehydrogenase. J Biol Chem 283, 9435–9443 (2008).

16. Jumper, J. et al. Highly accurate protein structure prediction with AlphaFold. Nature 596, 583–589 (2021).

17. Majorek, K. A. et al. The RNase H-like superfamily: new members, comparative structural analysis and evolutionary classification. Nucleic Acids Res 42, 4160–4179 (2014).

18. Prew, M. S. et al. Structural basis for defective membrane targeting of mutant enzyme in human VLCAD deficiency. Nat Commun 13, 3669 (2022).

19. Ashkenazy, H. et al. ConSurf 2016: an improved methodology to estimate and visualize evolutionary conservation in macromolecules. Nucleic Acids Res 44, W344–350 (2016).

20. West, A. P. et al. TLR signalling augments macrophage bactericidal activity through mitochondrial ROS. Nature 472, 476–480 (2011).

21. Hu, Y., Liu, W., Zhang, X. & Liu, D. ECSIT inhibits cell death to increase tumor progression and metastasis via p53 in human breast cancer. Transl Cancer Res 11, 699–709 (2022).

22. Canovas, B. & Nebreda, A. R. Diversity and versatility of p38 kinase signalling in health and disease. Nat Rev Mol Cell Biol 22, 346–366 (2021).

23. Honda, R., Korner, R. & Nigg, E. A. Exploring the functional interactions between Aurora B, INCENP, and survivin in mitosis. Mol Biol Cell 14, 3325–3341 (2003).

24. Pinho, C. M., Teixeira, P. F. & Glaser, E. Mitochondrial import and degradation of amyloid-beta peptide. Biochim Biophys Acta 1837, 1069–1074 (2014).

25. Kalaria, R. N. et al. Abundance of the longer A beta 42 in neocortical and cerebrovascular amyloid beta deposits in Swedish familial Alzheimer’s disease and Down’s syndrome. Neuroreport 7, 1377–1381 (1996).

26. Joh, Y. & Choi, W. S. Mitochondrial Complex I Inhibition Accelerates Amyloid Toxicity. Dev Reprod 21, 417–424 (2017).

27. Nsiah-Sefaa, A. & McKenzie, M. Combined defects in oxidative phosphorylation and fatty acid beta-oxidation in mitochondrial disease. Biosci Rep 36(2):e00313 (2016).

28. Lim, S. C. et al. Loss of the Mitochondrial Fatty Acid beta-Oxidation Protein Medium-Chain Acyl-Coenzyme A Dehydrogenase Disrupts Oxidative Phosphorylation Protein Complex Stability and Function. Sci Rep 8, 153 (2018).

29. Varadi, M. et al. AlphaFold Protein Structure Database: massively expanding the structural coverage of protein-sequence space with high-accuracy models. Nucleic Acids Res 50, D439–D444 (2022).

30. Miao, B., Xiao, Q., Chen, W., Li, Y. & Wang, Z. Evaluation of functionality for serine and threonine phosphorylation with different evolutionary ages in human and mouse. BMC Genomics 19, 431 (2018).

31. Juyoux, P., et al. Architecture of the MKK6-p38α complex defines the basis of MAPK specificity and activation. bioRxiv, doi:10.1101/2022.07.04.498667 (2022).

32. Kandiah, E. et al. CM01: a facility for cryo-electron microscopy at the European Synchrotron. Acta Crystallogr D Struct Biol 75, 528–535 (2019).

33. Punjani, A., Rubinstein, J. L., Fleet, D. J. & Brubaker, M. A. cryoSPARC: algorithms for rapid unsupervised cryo-EM structure determination. Nat Methods 14, 290–296 (2017).

34. Goddard, T. D. et al. UCSF ChimeraX: Meeting modern challenges in visualization and analysis. Protein Sci 27, 14–25 (2018).

35. Emsley, P., Lohkamp, B., Scott, W. G. & Cowtan, K. Features and development of Coot. Acta Crystallogr D Biol Crystallogr 66, 486–501 (2010).

36. Afonine, P. V. et al. Real-space refinement in PHENIX for cryo-EM and crystallography. Acta Crystallogr D Struct Biol 74, 531–544 (2018).

37. Mirdita, M. et al. ColabFold: making protein folding accessible to all. Nat Methods 19, 679–682 (2022).

38. UniProt, C. UniProt: the universal protein knowledgebase in 2021. Nucleic Acids Res 49, D480–D489 (2021).

39. Mullan, M. et al. A pathogenic mutation for probable Alzheimer’s disease in the APP gene at the N-terminus of beta-amyloid. Nat Genet 1, 345–347 (1992).

40. Spinazzi, M., Casarin, A., Pertegato, V., Salviati, L. & Angelini, C. Assessment of mitochondrial respiratory chain enzymatic activities on tissues and cultured cells. Nat Protoc 7, 1235–1246 (2012).

41. Shevchenko, A., Wilm, M., Vorm, O. & Mann, M. Mass spectrometric sequencing of proteins silver-stained polyacrylamide gels. Anal Chem 68, 850–858 (1996).

42. Cox, J. & Mann, M. MaxQuant enables high peptide identification rates, individualized p.p.b.-range mass accuracies and proteome-wide protein quantification. Nat Biotechnol 26, 1367–1372 (2008).

43. Schilling, B. et al. Platform-independent and label-free quantitation of proteomic data using MS1 extracted ion chromatograms in skyline: application to protein acetylation and phosphorylation. Mol Cell Proteomics 11, 202–214 (2012).

